# The functional form of specialized predation dramatically affects whether Janzen-Connell effects can prevent competitive exclusion

**DOI:** 10.1101/2021.07.10.451901

**Authors:** Daniel J. B. Smith

**Author notes:** Corresponding Author: Daniel J. B. Smith.

## Abstract

Janzen Connell Effects (JCEs), specialized predation of seeds and seedlings near conspecific trees, are hypothesized to promote high species richness. While past modeling studies show JCEs can maintain higher diversity than a neutral community, recent theoretical work indicates JCEs may weakly inhibit competitive exclusion when species exhibit interspecific fitness variation. However, recent models make somewhat restrictive assumptions about the functional form of specialized predation – that JCEs occur at a fixed rate when seeds/seedlings are within a fixed distance of a conspecific tree. Using a theoretical model, I show that the functional form of JCEs largely impacts their ability to promote coexistence. If specialized predation pressure increases additively with adult tree density and decays exponentially with distance, JCEs maintain considerably higher diversity than predicted by recent models. Parameterizing the model with values from a Panamanian tree community indicates JCEs can maintain high diversity in communities exhibiting high interspecific fitness variation.

## Introduction

The Janzen-Connell Hypothesis (JCH) is a long-studied species coexistence mechanism frequently invoked to explain the high diversity of tropical forests (Janzen, 1970; Connell, 1971; Wright, 2002; Terborgh, 2012, 2020). The JCH is based on the observation that specialized natural enemies (predators, fungi, and pathogens) attack seeds and seedlings near conspecific trees. These phenomena are frequently referred to as Janzen-Connell Effects (JCEs). JCEs are thought to promote coexistence by creating negative frequency dependence – as a species becomes more common, its seeds and seedling experience greater predation pressure. Rare species, experiencing less predation pressure, enjoy a fitness advantage. This is thought to prevent extinction and maintain diversity. However, the efficacy of this mechanism remains contested on theoretical grounds.

The conditions necessary for the JCH to operate, the presence of distant or density-dependent specialized natural enemies that decrease seed and seedling survivorship, enjoy empirical support in a variety of systems (e.g. Hyatt *et al.*, 2003; Petermann *et al.*, 2008; Mangan *et al.*, 2010; Swamy & Terborgh, 2010; Zhu *et al.*, 2010; Johnson *et al.*, 2012, 2014; Comita *et al.*, 2014; Bever *et al.*, 2015; Jevon *et al.*, 2020; Hazelwood *et al.*, 2021). Additionally, theoretical work demonstrates that JCEs effectively prevent extinction due to ecological drift and maintain higher species richness than neutral models (Armstrong, 1989; Adler & Muller-Landau, 2005; Sedio & Ostling, 2013; Levi *et al.*, 2019). However, a coexistence mechanism must be able stabilize coexistence in the presence of interspecific fitness differences (Chesson, 2000). Recent studies that integrate fitness differences into JCE models call into question their to promote deterministic coexistence (Hülsmann *et al.*, 2020; Cannon *et al.*, 2021). Most relevantly, models that consider inter-specific variation in intrinsic fitness indicate that JCEs may be unable to reduce competitive differences to the point of community-wide coexistence, causing some to label JCEs as “a weak impediment to competitive exclusion” (Chisholm & Fung, 2020).

A notable assumption of several recent JCH-type models (e.g. Levi *et al.*, 2019; Chisholm & Fung, 2020) is the functional form of specialized predation pressure. These studies assume that JCEs affect seeds and seedlings within a fixed distance (or “neighborhood”) of a conspecific adult tree. Where JCEs affect seeds and seedlings, they reduce survivorship by a fixed proportion that is independent of conspecific adult density. I refer to this as the “fixed neighborhood model” (Fig. 1A and 1D). This model is a reasonable assumption if JCEs are primarily the consequence of qualitative modifications to soil composition, an outbreak of a pathogen that occurs at a threshold seed or seedling density, or other static ways in which the presence of a species might modify the local environment. An alternative JCE functional form is one in which seed and seedling mortality increases additively with conspecific adult density or basal area and declines monotonically with conspecific distance. I refer to this as the “additive predation model” (Fig. 1C, 1F). Several studies use this functional form (e.g. Adler & Muller-Landau, 2005; Muller-Landau & Adler, 2007; Sedio & Ostling, 2013; Stump & Comita, 2020; Krishnadas & Stump, 2021). If seed/seedling mortality increases with the local abundance of natural enemies that disperse from nearby trees, this JCE model is more realistic. Indeed, JCE strength is often quantified by examining how a species’ density at one life history stage decreases its survivorship at another stage (e.g. Harms *et al.*, 2000; Klironomos, 2002; Mangan *et al.*, 2010; Johnson *et al.*, 2012; Bagchi *et al.*, 2014; LaManna *et al.*, 2016; Fricke & Wright, 2017) and empirical evidence indicates the additive predation model reflects nature, at least is some communities (Comita *et al.*, 2010; Liu *et al.*, 2015). Previous theory does not examine the difference between these two JCE functional forms and it is unknown how they might differently impact highly diverse communities that vary in fitness.

**Fig. 1.**
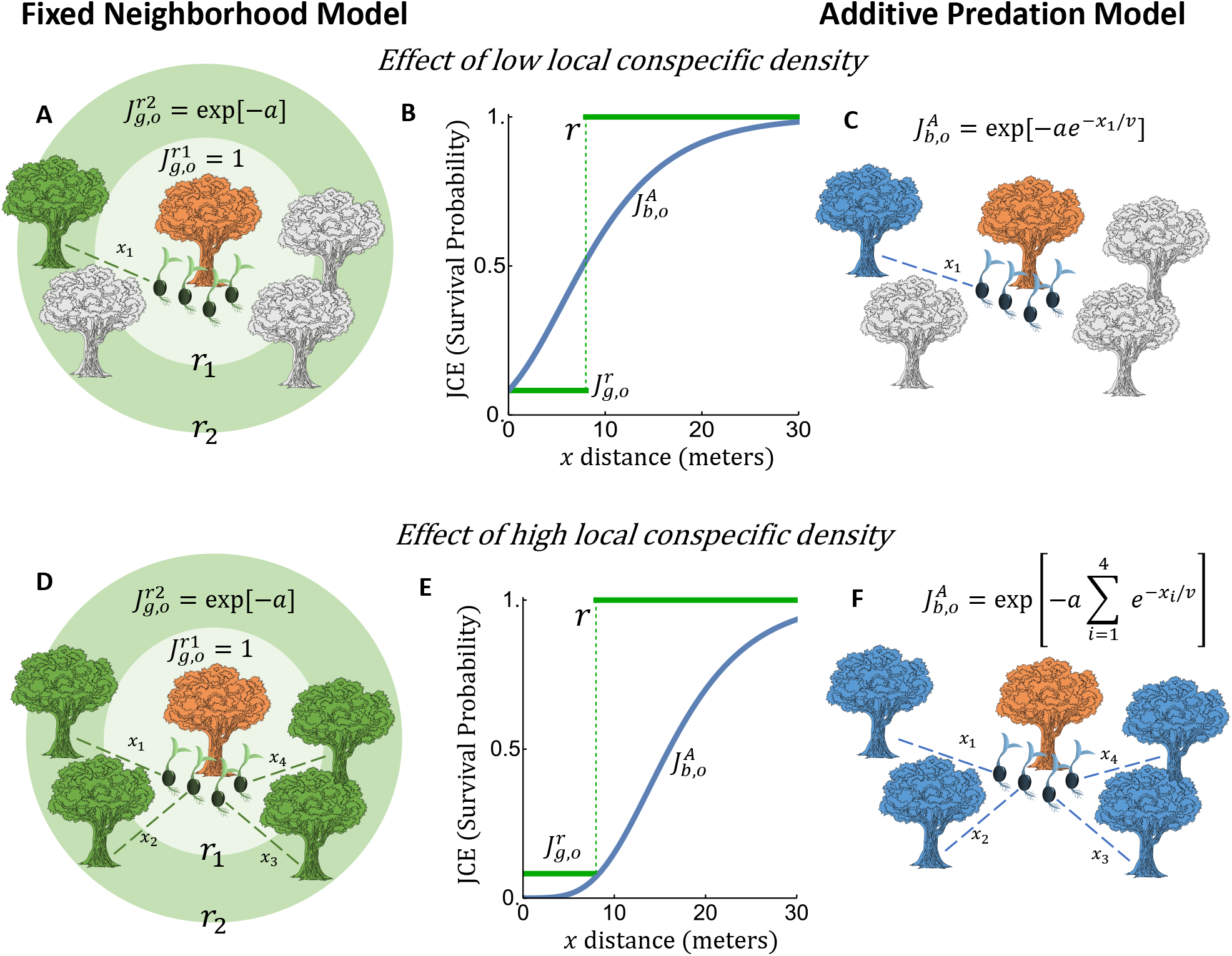
Visualization of the specialized predation functional forms (A, C, D, and F) and their implications for seedling survivorship at low and high conspecific densities (B and E). Panels A and D depict the fixed neighborhood model. Each panel depicts the impact conspecific adult density (green trees) on seedling survivorship (green seedlings) on a focal patch occupied by a heterospecific species (the orange tree). Grey trees depict unspecified heterospecifics. In panels A and D, two radii (*r*_1_ and *r*_2_) depict two different fixed neighborhood effect scenarios. In each panel, if no conspecific (green) tree falls within the radius, then its seedlings on the focal patch do not experience JCEs. This occurs when *r*_1_ defines the neighborhood effect (*x*_1_ or min(*x*_1_*, x*_2_*, x*_3_*, x*_4_) > *r*_1_). In this case, 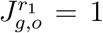 where 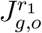 is the survivorship of the green (*g*) species’ seedlings on the orange (*o*) patch when the neighborhood effect is of radius *r*_1_. When the radius is *r*_2_, conspecific trees fall within the JCE neighborhood (*x*_1_ or min(*x*_1_*, x*_2_*, x*_3_*, x*_4_) < *r*_1_) and the seedlings experience the fixed JCE: 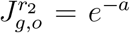. Notably, in panel D, seedlings do not experience JCEs if the distance is fixed at *r*_1_ despite the high density of conspecifics in the local area. For the *r*_2_ case in panel D, multiple trees falling in the exclusion zone does not increase the strength of the JCE (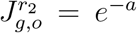 in both panel A and panel D). Panels C and F depict the additive predation model. Each panel depicts the impact conspecific density (blue trees) on seedling survivorship (blue seedlings) on a focal patch occupied by a heterospecific species (orange). Grey trees depict unspecified heterospecific trees. In panel C, a single conspecific (blue) tree is near the focal orange patch and a portion of 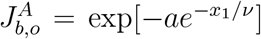 blue seedling survive where *x*_1_ is the distance of the conspecific tree from the focal patch. Panel F depicts four conspecific (blue) trees near the focal patch; blue seedling survivorship is an additive function of all the nearby conspecifics scaled by distance: 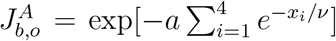. Panels B and E depict the survivorship of seedlings based on proximity to conspecific trees as defined by each JCE functional form. The *y*-axis is seedling survivorship and the *x*-axis is conspecific distance(s) from the focal patch. The green lines depict the fixed neighborhood model and the blue curves depict the additive predation model. Panel B shows the case of a single nearby conspecific (as depicted in panels A and C). The conspecific is *x* meters away from the focal patch. For the fixed neighborhood model, seedling survival is *e^−a^* if the conspecific falls within the radius, *r*, and 1 otherwise. For the additive predation model, seedling survivorship increases monotonically with distance. The additive predation model induces less mortality than the fixed neighborhood model if 0 < *x* < *r* and greater mortality if *x* > *r*. In panel E, it is assumed that the focal patch is surrounded by 4 conspecifics (as depicted in panels D and F). For simplicity, all conspecifics are equidistant from the focal patch (*x* = *x*_1_ = *x*_2_ = *x*_3_ = *x*_4_). For the fixed neighborhood model, seedling mortality behaves identically to the single tree case with respect to *x*. For the additive predation model, seedling surviorship is lower than that of the single conspecific case for all *x*. In addition, seedling survivorship in the additive predation model is always lower than that of the fixed neighborhood model. In each plot, *a* = 2.5, *r* = 8.0, and *ν* = 6.0.

**Table 1.** Relevant variables, terms, parameters, and parameter ranges considered. Cited parameter ranges are based on empirical studies or ranges considered in previous models.

| Parameter/<br>Variable | Description | Estimated value/<br>range considered | Reference (if relevant) |
| --- | --- | --- | --- |
| $p_i$ | Proportion of species $i$ in the community | - | - |
| $N$ | Community diversity | 300 | - |
| $Y_i$ | Intrinsic fitness of species $i$ | $Y \sim \text{lognormal}[0, \sigma_Y]$ | - |
| $\sigma_Y$ | Standard deviation of $Y$ | 0.1 – 1.0 | Chisholm & Fung (2020) |
| $a_i$ | Specialized predation pressure of species $i$ | 0.1 – 5 | - |
| $d$ | Proportion of seeds dispersed | 0.1 – 1.0 | - |
| $J_{i,i}(p_i)$ | Local JCE functional form | - | - |
| $J_{i,k}(p_i)$ | Non-local JCE functional form | - | - |
| $r$ | Neighborhood radius of specialized predation | - | - |
| $\gamma$ | Tree density (individuals per square meter) | 0.17 – 0.42 | Condit *et al.* (2019) |
| $\nu$ | Distant decay rate of specialized predation | 5 – 10 | Comita *et al.* (2010) |
| $E_n$ | Neighborhood effect parameter of the fixed neighborhood model; $\pi\gamma r^2$ | 8, 24, 49 | Chisholm & Fung (2020)<br>Levi *et al.* (2019) |
| $E_A$ | Neighborhood effect parameter of the additive model; $2\pi\gamma\nu^2$ | $\approx 27 - 125$ | From constituent parameters assuming $\gamma = 0.20$ |

In this paper, I develop Ordinary Differential Equation (ODE) approximations of spatially explicit JCE models. I use the ODEs to compare the relative ability of each JCE functional form to promote deterministic coexistence. I find that the additive predation model promotes considerably greater diversity than the fixed neighborhood model when species experience inter-specific variation in intrinsic fitness. I conclude that JCEs are capable of maintaining high species richness if they follow the additive predation model and are sufficiently strong relative to the magnitude of interspecific fitness variation. This study highlights the need to empirically determine the functional form of distance dependent predation and quantify the key parameters that affect its strength.

## Model and analysis

Modeling methods are similar to Chisholm & Muller-Landau (2011), Levi *et al.* (2019), and Chisholm & Fung (2020). I consider a tree community of *N* species that contains *M* patches in which every patch contains a single adult tree. Trees disperse seeds across patches which grow into seedlings. On each patch, seedlings experience JCEs based on their proximity to conspecific trees (for simplicity, I assume JCEs affect seedlings). When an adult tree dies, a seedling is chosen at random to replace it (a lottery model; Chesson & Warner (1981)).

### Seed and seedling dynamics

Seedling abundances on each patch are determined as follows: (1) each tree produces seeds at a constant rate. All trees uniformly disperse a portion of their seeds (*d*) among patches and retain the remaining portion of their seeds (1 − *d*) on the local patch. (2) *Y_i_*, henceforth intrinsic fitness, defines the effective number of seedlings species *i* produces. *Y_i_* is a composite parameter of seed production rate, baseline seed and seedling survival rate, and competitive ability. I assume dispersed seeds develop into seedlings upon competition for a patch. (3) JCEs kill a portion of seedlings on each patch based on their proximity to conspecific trees through either the fixed neighborhood or additive predation model. Let *S_x,y_* be the abundance of seedlings of species *x* on a patch occupied by species *y*. Seedling abundances are:

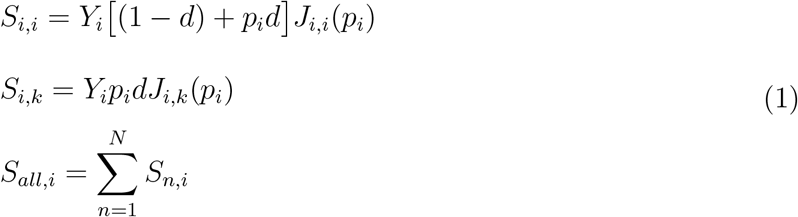

where *p_i_* is the proportion of species *i* in the population. *J_i,i_*(*p_i_*) and *J_i,k_*(*p_i_*) are functions that depict the JCE-induced seedling survivorship of species *i* on a patch occupied by species *i* and *k*, respectively (*i* ≠ *k*). These functions contains spatial information. I assume each species’ seedlings are at the above values upon the death of the adult in the patch.

### Fixed neighborhood model

Similar to Levi *et al.* (2019) and Chisholm & Fung (2020), I consider when JCEs kill a fixed proportion of a species’ seedlings on patches within *r* meters of a conspecific adult (Fig. 1A and 1D). Let *E_n_* adjacent patches fall within the neighborhood set by *r*. The seedling survivorship functions for species *i* under the fixed neighborhood model are:

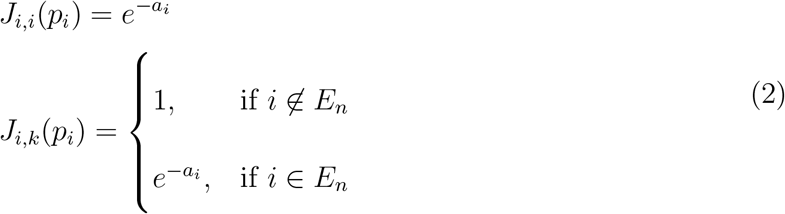

for *k* ≠ *i*. *i* ∈ *E_n_* is the condition that a conspecific tree of species *i* is inside the neighborhood (radius) of a patch occupied by a heterospecific tree (*k*) and *i* ∉ *E_n_* is when *i* is not within the neighborhood. *a_i_* is a composite parameter of seedling predation rate and the time over which seedling predation occurs. I assume seedlings experience specialized mortality at rate 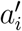 over a period of time *τ*. Therefore, if *Y_i_d* seedlings would be present in the absence of JCEs, then 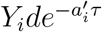 survive predation. Hence, 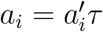.

### Additive predation model

I consider when seedlings experience predation pressure that increases linearly as a function of local conspecific density and decreases exponentially with distance (Fig. 1C and 1F). The predation pressure experienced by species *i*’s seedlings located *x* meters away from a single conspecific adult is *a_i_e^−x/ν^* where *ν* defines rate at which predation declines with distance. *a_i_e^−x/ν^ = a_i_* when *x* = 0, in which case *a_i_* can be interpreted as the maximum predation pressure seedlings can experience from a single conspecific adult. Incorporating additive predation yields the following seedling survivorship functions:

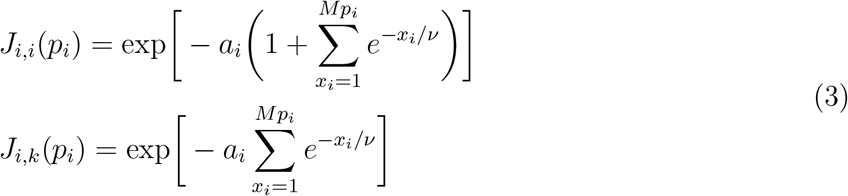

where *x_i_* is the distance of the *x_th_* closest individual of species *i* to the focal patch and *Mp_i_* is the number of individuals of species *i* in the population. *J_i,i_*(*p_i_*) contains the +1 term because the local patch contains a conspecific adult. Seedling are assumed to be at the center of the each patch.

### Tree dynamics

All adult trees die at rate *δ*. When a tree dies, it is immediately replaced by a randomly selected seedling in the patch. Let *P* (*x, y*) be the probability a seedling of species *x* colonizes a patch previously occupied by species *y*. *P* (*i, i*) = *S_i,i_/S_all,i_* and *P* (*i, k*) = *S_i,k_/S_all,k_*. These quantities can be implemented into a Spatially Explicit Model (SEM). Alternatively, the dynamics can be deterministically approximated by the expected seedling abundances of each patch type and implementing them into the following system of ODEs:

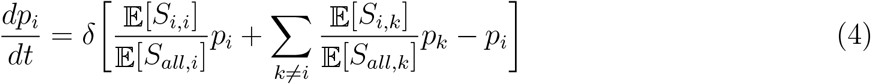

for *i* = 1, 2, …, *N*. The first term in the parentheses is the rate at which species *i* recolonizes a patch previously occupied by a conspecific, the second term is the rate at which species *i* colonizes a patch previously occupied by species *k* (*k* ≠ *i*), and the third term is the rate atwhich patches occupied by species *i* are disturbed. The ODE model approximates the behavior of the SEM (Appendices A and B, “SEM and derivation of ODE approximation”).

Under the ODE, the fixed neighborhood model seedling equations are:

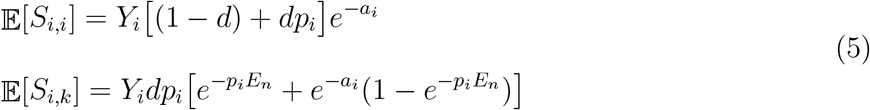

where *E_n_* = *γπr*^2^. *γ* is tree density (trees per square meter). *E_n_* is the expected number of trees within a radius of *r* meters. If the neighborhood effect extends *r* meters, there will be on average *γπr*^2^ individuals within the neighborhood. See Appendix A for details. For the additive predation model, the seedling equations are:

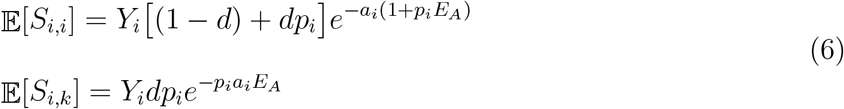

where *E_A_* = 2*πγν*^2^. *p_i_a_i_E_A_* is the expected predation pressure experienced by species *i* (see Appendix B for details). 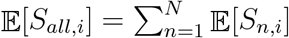 in both cases.

### Parameterizations

I roughly parameterized key quantities of the model using data from the Barro Colordo Island (BCI) forest plot in Panama. Tree density (*γ*) can be approximated by dividing the total number of individuals in the community by its area in square meters. The plot at BCI is 50-ha and contains approximately 210, 000 live trees (Condit *et al.*, 2019). However, as Chisholm & Fung (2020) note, there are approximately 86, 000 trees of reproductive height. Distant-dependent predation pressure may emerge from only adult trees or could manifest in earlier stages of tree development. Therefore, to a first approximation, the appropriate number of individuals to calculate *γ* is likely in between these values. This yields a range of *γ* between 0.17 and 0.42 individuals per square meter. *ν*, the rate at which predation declines as a function of distance in meters, was estimated in Comita *et al.* (2010) with best and second best fit values equivalent to *ν* = 5 and *ν* = 10, respectively. I considered this range of *ν* values.

### Presentation of results

#### ODE analysis

I compared the relative ability of each JCE functional form to promote coexistence in communities experiencing interspecific variation in intrinsic fitness (*Y*) with equal JCE susceptibility. I derived an approximate invasion criteria for each model. I then performed ODE simulations to quantify the species richness each JCE functional form maintains. I considered inter-specific variation in *Y* to be log-normally distributed: *Y* ~ lognormal[*μ* = 0*, σ_Y_*]. I explored values of *σ_Y_* between 0.1 − 1.0 and values of *a* between 0.1 − 5.0. For the fixed neighborhood model, I ran simulations with *E_n_* = 8, 24, and 49. These values were selected for consistency with previous models (Hubbell, 1980; Levi *et al.*, 2019; Chisholm & Fung, 2020) and correspond to when JCEs occur in the 3×3, 5×5, and 7×7 Moore neighborhoods surrounding a focal tree/patch. For the additive predation model, I examined *ν* = 5, 7.5, and 10. I performed simulations for *d* ~ {1.0, 0.6, 0.1}.

All simulations began with 300 species of equal frequency in the population. 300 was selected on the basis that BCI contains approximately 300 woody plant species (Condit *et al.*, 2019). ODE simulations were run for 6,250 generations, far longer than sufficient for the system to reach equilibrium. I considered species *i* to be extinct if log(*p_i_*) < −11, where log(*p_i_*) = −11 corresponds to a species with an expected abundance of approximately one adult tree at BCI (log(1/86000) ≈ −11.4). I assumed *γ* = 0.20. Simulations were performed in R (R Core Team, 2020).

#### ODE validation

To validate the use of the ODE model, I ran SEM and ODE simulations using identical parameters and compared outputs in terms of species richness and effective species richness (exp(Shannon Diversity)). SEM simulations were run until the transient dynamics had approximately concluded. I compared the SEM to a slightly modified version of the ODE model that takes into account the structure of the SEM. Simulations were conducted over a parameter range similar to the above-listed ODE simulations. I ran simulations with *σ_Y_* ~ {0.1, 0.45, 0.8} and *a* ~ {1, 2.75, 4.5} with *d* = 1. For each of the 9 parameter combinations, I ran simulations with *E_n_* ~ {8, 24, 49} for the fixed neighborhood model and *ν* ~ {5, 7.5, 10} for the additive predation model. See appendices A and B for details.

## Results

### ODE validation

The SEM and ODE model produced similar outputs of species richness and effective species richness (Fig. 2). While inexact, this indicates the ODE models to be suitable first order approximations of the SEMs (see “Results of comparison”, Appendices A and B).

**Fig. 2.**
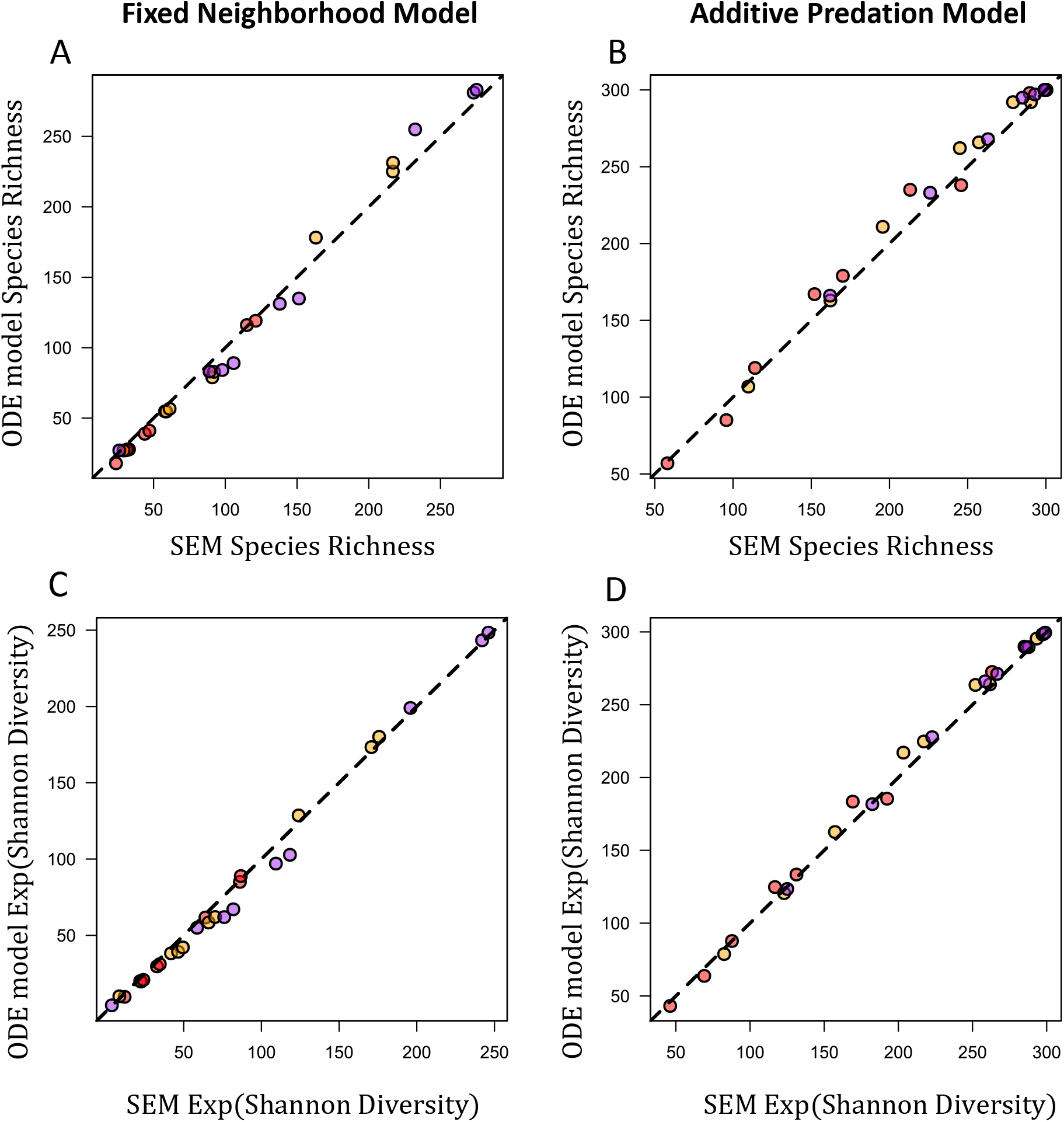
ODE model validation. Panels compare the species richness and the effective species richness (the exponential of Shannon Diversity) from outputs of the SEM and the ODE model under identical parameterizations. Panels A and C show the fixed neighborhood model and panels B and D show the additive predation model. The dashed line is the one-to-one line (points on the line represent when the SEM and ODE yield the exact same diversity output). Red points are when *ν* = 5 or *E_n_* = 8, orange/yellow points are when *ν* = 7.5 or *E_n_* = 24, and purple points are when *ν* = 10 or *E_n_* = 49. The ODE model yields outputs highly similar the SEM by both diversity metrics for both models.

### ODE analysis

#### Invasion criteria

A species can deterministically invade a community if it has a positive invader per capita growth rate (Turelli, 1978; Chesson, 2000). Therefore, comparing the invader per capita growth rate of each JCE functional form provides a metric of their relative ability to promote coexistence. Assuming species vary only in intrinsic fitness (*Y*) and *d* = 1, the invasion criteria of species *i* with intrinsic fitness *Y_i_* invading a community of *N* resident species is approximately

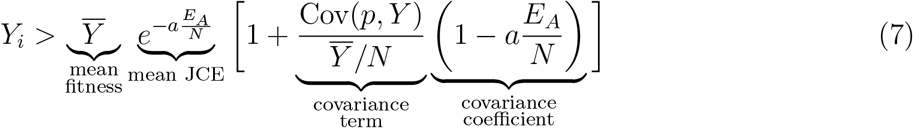

for the additive predation model and

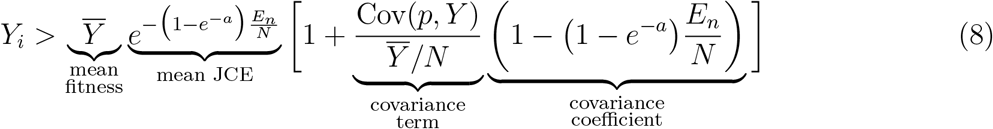

for the fixed neighborhood model (similar to the derivation in Chisholm & Fung (2020)). The right hand side of each invasion criteria indicates the minimum invader fitness that permits invasion. See “Derivation of invasion criteria” in Appendices A and B for details.

The right hand side of each invasion criteria is broken into four parts. (1) Mean fitness is the average intrinsic fitness of the resident community. Invasion becomes more difficult with increasing 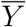. (2) The mean JCE is the average proportion of seedlings that survive JCEs in the resident community. This defines the average fitness-equalizing strength of each model. The smaller the mean JCE, the lower the mean seedling survivorship and the easier invasion is. (3) The covariance term represents the relationship between species proportion and intrinsic fitness (if it is positive, species with higher fitness make up a relatively high proportion of the community). Invasion becomes more difficult as the covariance term increases – the higher the covariance, the greater the proportion of high fitness species the invader must compete against. (4) The covariance coefficient relates to how JCEs disproportionately affect common species with higher fitness. If species with high intrinsic fitness make up a relatively large proportion of the community, their seedlings experience relatively high JCE-induced mortality. This reduces the negative effect of the covariance term. The smaller the covariance coefficient is, the less competition the invader experiences from high fitness competitors and the easier invasion becomes.

Examination of the mean JCE term yields some insights. *a* differently impacts each mean JCE term, reflecting how each functional form produces negative frequency dependence. The fixed neighborhood model sets an upper bound to the strength of JCEs (*e^−a^*) over a fixed area, *E*. The mean JCE term in equation (8), 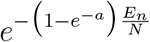, is a saturating function of *a*: for large *a*, it converges to 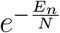 in which all conspecific seedlings of the species found within *E_n_* die. 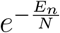 reflects the average probability a resident species experiences JCEs. If *N* is large relative to *E_n_*, species are very unlikely to be found in the neighborhood. Frequency dependence is thus strictly limited by *E_n_*, which must be very large for JCEs to equalize fitness in diverse communities, even if *a* is arbitrarily large. In contrast, the additive predation model has neither a fixed bound on the strength of JCEs nor the distance over which they occur. Instead, seedling predation decays exponentially with distance and increases with local conspecific density. The mean JCE term in equation (7), 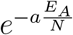, converges to zero as *a* becomes large (meaning that any species with positive fitness can invade). Underlying this result is the interaction between additive and exponentially decaying predation pressure. Because of the exponential decay functional form, relatively weak JCEs affect seedlings when a single conspecific tree is a moderate distance away (Fig. 1B, 1C). Because predation pressure acts additively, moderate distance JCEs become fairly strong when a species is common (Fig. 1E, 1F). As predation pressure is proportional to *a* at all distances, the strength of frequency dependence is also proportional to *a*. The same arguments apply for covariance coefficients of each model, which contain the same expressions.

An illuminating quantity summarizing the relative ability of each model to promote coexistence is the ratio of their invasion criteria (equation (8) divided by equation (7), henceforth Φ):

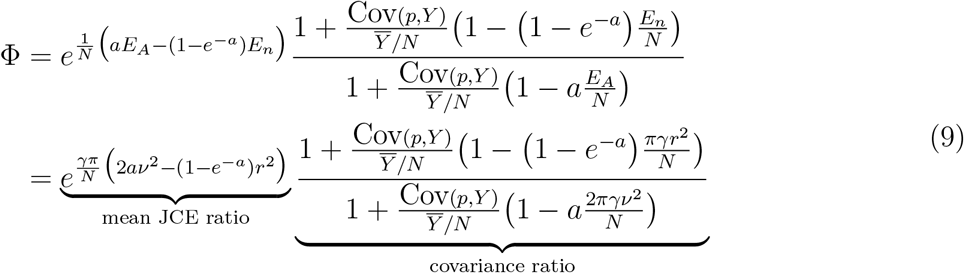

Φ can be interpreted in the following way. Given a set of parameters, let 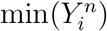 and 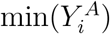 be the minimum intrinsic fitness that allows species *i* to invade the resident community under the fixed neighborhood model and the additive predation model, respectively. 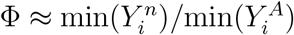. If Φ > 1, the additive predation model reduces fitness differences more effectively than the fixed neighborhood model. Emphasizing the mean JCE ratio, examination of Φ has several immediate implications. The additive model becomes increasingly more effective as *a* increases; Φ → ∞ as *a* → ∞ (Fig. 3A, 3C). This is consistent with the above analysis. Second, Φ increases as a function of *γ/N*, meaning the additive predation model disproportionately reduces fitness differences in communities of high average tree density (Fig. 3B, 3D). Third, Φ increases with the spatial scale of distant-dependent predation. Assuming *ν* = *r*, larger *ν* yields much greater values of Φ (Fig. 3).

**Fig. 3.**
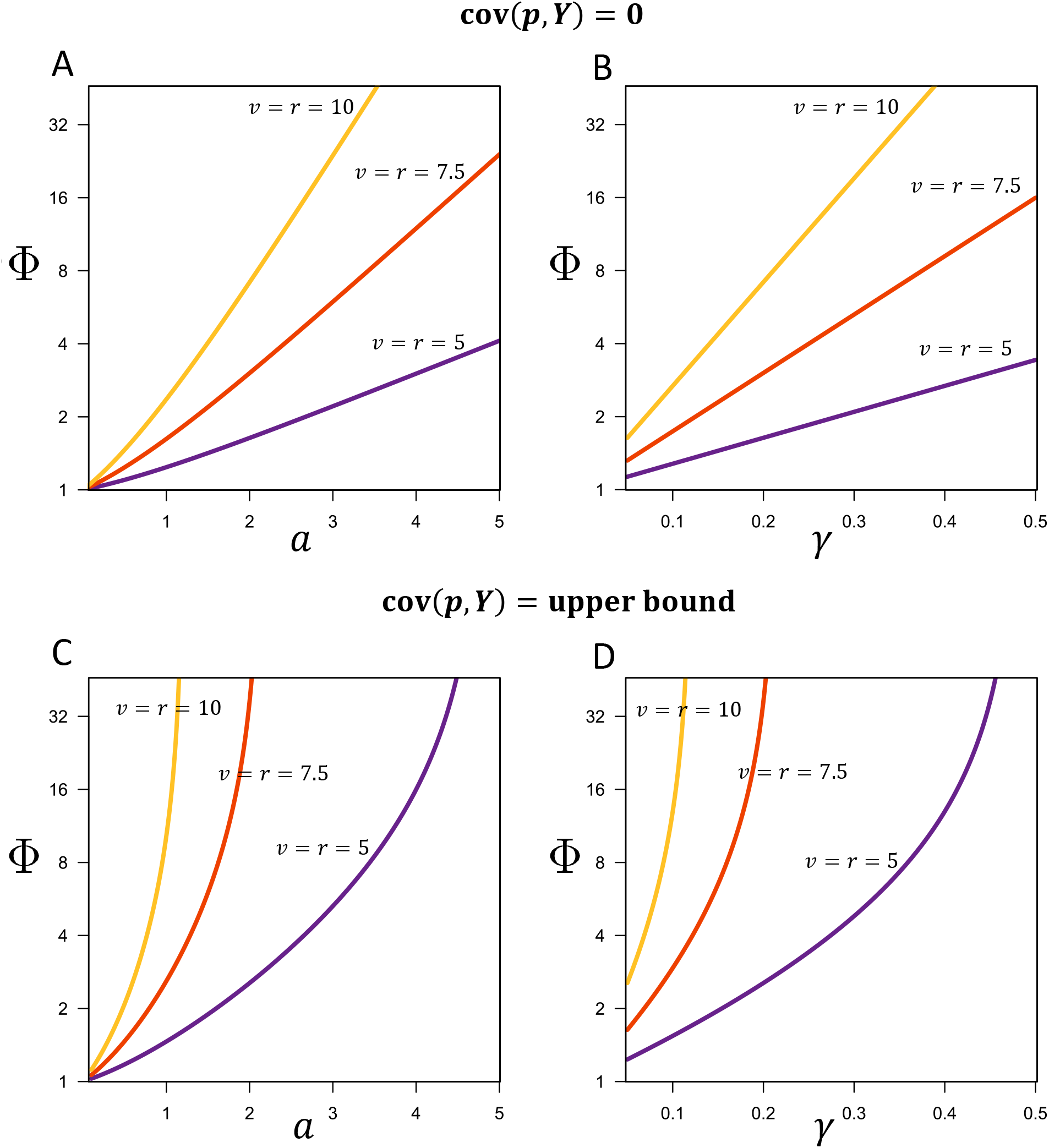
The impact of different parameters on Φ (the relative efficacy of the fixed neighborhood model and the additive predation model in reducing fitness differences). Φ > 1 indicates the additive predation model more effectively reduces fitness differences by a factor of Φ. For all panels, the *y*-axis is Φ. Note the log scale on the *y*-axis. Panels A and B show when cov(*p, Y*) = 0 and panels C and D show when 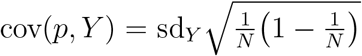 (the upper bound). Panels A and C show the effect of *a* (specialized predation rate) on Φ for *γ* = 0.2. For all *a* tested (0.1 − 5.0), the additive predation model more effectively reduces fitness differences (Φ > 1). This effect is more significant when predation pressure extends over a larger distance (compare the three lines, as indexed by *ν* and *r* values). The values of *E_n_* correspond to 15.7, 35.3, and 62.8 (*E_n_* = *γπr*^2^). Panels B and D show the effect of *γ* (tree density) on Φ with *a* = 2.0. Similar to the above case, greater *γ* corresponds to higher Φ. Overall, the additive predation model is particularly effective when predation is strong (large *a*), trees are of high density (large *γ*), and the spatial scale of predation is large (large *ν* and *r*). These effects are stronger when cov(*p, Y*) is large. In panels C and D, *σ_Y_* = 0.5. For both panels, *N* = 100.

I numerically examined Φ under lower and upper bounds of Cov(*p, Y*). First, I examined the lower bound Cov(*p, Y*) = 0, in which the covariance ratio converges to 1 and the expression collapses the mean JCE ratio. Second, I examined when the covariance is at its upper bound: 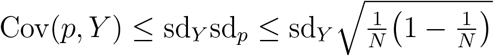 where sd*_Y_* and sd*_p_* are the standard deviations of intrinsic fitness and species proportion, respectively. The upper bound of sd*_p_*, 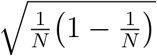, was derived using the Bhatia–Davis inequality (Bhatia & Davis, 2000). Each case shows the same qualitative pattern: the additive predation model tends to more strongly reduce fitness differences (Fig. 3). However, the discrepancy is considerably greater when the covariance term is high (compare Fig. 3A and 3B to 3C and 3D). This reflects the greater frequency dependence the additive predation model induces. See Appendix C for a validation of the accuracy of Φ.

#### ODE simulations

ODE simulations indicate that the additive predation model promotes higher levels of species richness than the fixed neighborhood model, consistent with the invasion analysis. For the fixed neighborhood model, species richness increased with *E_n_* and increased as a saturating function of *a* (Fig. 4A, 4C, 4E). Similar to Chisholm & Fung (2020), the fixed neighborhood model maintained high species richness only under low inter-specific variation in intrinsic fitness: species richness was relatively high for *σ_Y_* ≈ 0.1, but eroded rapidly as *σ_Y_* increased for all *E_n_*. In contrast, the additive predation model promoted high species richness for all *ν* (5, 7.5, and 10) especially the latter two values (Figs. 4B, 4D, 4F). Species richness increased approximately linearly with *a*. Species richness was high unless *a* was relatively low and *σ_Y_* was high. For *ν* = 10, community-wide coexistence was possible, even with *σ_Y_* = 1. Similar to (Stump & Chesson, 2015) and Chisholm & Fung (2020), greater dispersal limitation (*d* = 0.6 or *d* = 0.1) slightly reduced diversity in each model, but did not qualitatively change results (Appendix E, Fig. E1-E2). Results showed the same qualitative pattern for Shannon Diversity (Appendix E, Fig. E3). Larger values of *E_n_* (higher *γ* or *r*) only marginally increased the diversity maintained by the fixed neighborhood model, but larger values of *E_A_* (higher *γ* or *ν*) dramatically increased the diversity maintained by the additive predation model (Appendix E, Fig. E4).

**Fig. 4.**
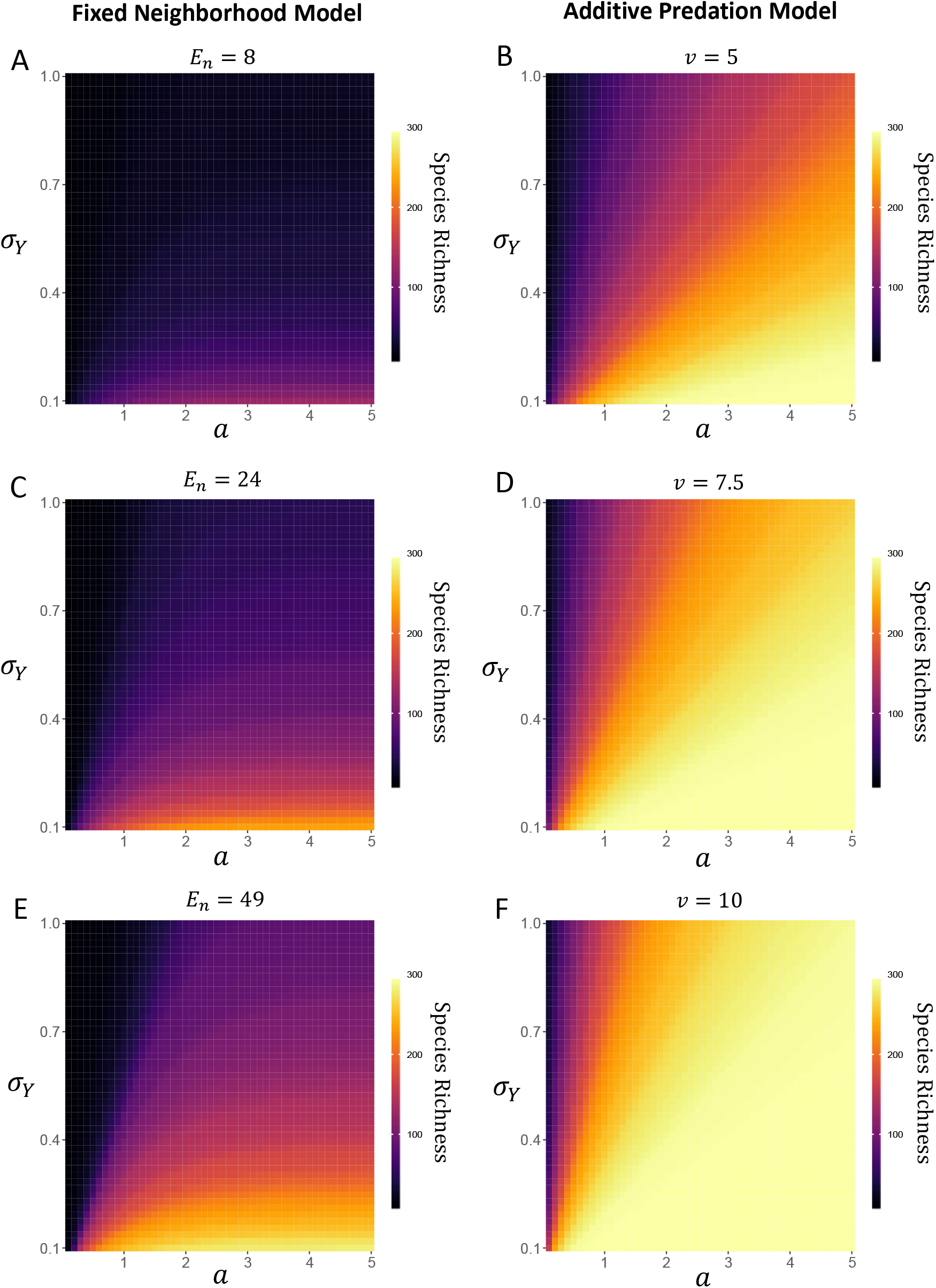
Species richness from ODE model outputs. For each heatmap, the *x*-axis is *a*, the *y*-axis is *σ_Y_* (*Y* ~ lognormal[*μ* = 0, *σ_Y_*]), and the *z*-axis is species richness. Each heatmap is composed of 2500 ODE outputs. Panels A, C, and E depict the neighborhood effect model for *E_n_* = 8, 24, and 49, respectively. Panels B and D depict the additive predation model when *ν* = 5, 7.5, and 10, respectively. Under these parameterizations, the additive predation model tends to maintain much higher species richness. *d* = 1 and *γ* = 0.2 in all simulations.

## Discussion

The Janzen-Connell Hypothesis (JCH), historically favored as a likely candidate to promote high species richness in tropical forests (Janzen, 1970; Connell, 1971; Wright, 2002; Terborgh, 2012) has recently been the target of harsh criticisms (Chisholm & Fung, 2020; Hülsmann *et al.*, 2020; May *et al.*, 2020; Cannon *et al.*, 2021). Theoretical works integrating fitness differences into JCH-type models argue that JCEs may be unable to promote high levels of diversity. This is a valid critique: for a coexistence mechanism to function, it must mitigate interspecific fitness differences (Chesson, 2000). In this paper, I assess how the functional form of specialized predation pressure affects the ability of JCEs to inhibit competitive exclusion in diverse communities by using ODE approximations of spatially explicit JCE models.

Recent theoretical studies (e.g. Levi *et al.*, 2019; Chisholm & Fung, 2020) assume that specialized predation pressure affects seeds and seedlings at a fixed rate within a predefined distance of a conspecific tree (the fixed neighborhood model). I compared the fixed neighborhood model to when specialized predation pressure increases as an additive function of conspecific density and declines exponentially with conspecific distance (the additive predation model). When species vary inter-specifically in fitness, the additive predation model promotes considerably higher diversity (Fig. 4). The discrepancy in species richness maintained by each JCE functional form stems from the distinct types of negative frequency dependence they generate. For the fixed neighborhood model, as it becomes more common, a larger proportion of patches in the community fall within a species’ JCE neighborhood (*E_n_*). However, the strength of frequency dependence is strictly limited by the fixed JCE strength and the size of the neighborhood. For the additive predation model, JCE strength increases with a species’ frequency in the population. This promotes a stronger advantage to rare species: their offspring are both less likely to experience JCEs than common species and experience weaker JCEs where they do. Additionally, the exponential decay of predation pressure permits JCEs to facilitate mortality at moderate distances. This interacts with additive predation to promote strong JCEs for relatively common species community-wide. This is consistent with the statistical metric proposed by Fricke & Wright (2017) which emphasizes how JCEs disproportionately affect the demography of common species. I also find that the additive predation model strongly reduces diversity when tree density is high (Fig. 3B, 3D), consistent with the hypothesis that JCEs are particularly important in dense tropical forest communities.

A potential limitation of this model is its treatment of seed dispersal. I assumed trees retain a proportion of seeds (1 – *d*) on the local patch and evenly distribute the remainder (*d*) throughout the community. Realistic dispersal kernels influence the spatial distribution of trees, making them more aggregated in space than expected by chance. This can influence competitive outcomes (Detto & Muller-Landau, 2016; Stump & Comita, 2020; Wiegand *et al.*, 2021). This issue is likely unimportant if the spatial scale of seed dispersal is sufficiently greater than that of seed/seedling predation. Chisholm & Fung (2020) found that model outputs using a dispersal kernel similar to fitted values from BCI data were almost identical to the global dispersal case for the fixed neighborhood model. The same is likely true for the additive predation model, although factors such as inter-specific variation in dispersal ability (e.g. Muller-Landau *et al.*, 2008) are also likely to modify results. Overall, while the treatment of dispersal in this study ignores a great deal of biology and may quantitatively influence results, it is unlikely to qualitatively affect the main conclusion that the additive predation model promotes greater diversity than the fixed neighborhood model.

There are several other caveats. First, I only examined inter-specific variation in intrinsic fitness, but species also vary in JCE susceptibility. Variation in JCE susceptibility in combination with dispersal limitation can induce competitive exclusion (Stump & Comita, 2018). Second, this paper emphasizes deterministic coexistence via analyzing ODEs, which ignores the potential effects of drift. Although JCEs can stabilize against drift in an otherwise neutral community (Levi *et al.*, 2019), inter-specific variation in fitness induces variance in species’ equilibrium abundances and weaker competitors may be more susceptible to stochastic extinction (Nisbet & Gurney, 1982; Miranda *et al.*, 2015). This should be explored for the additive predation model in future studies. Third, this paper does not consider species immigration. The model presented by May *et al.* (2020) suggests immigration could drown out the signal of JCEs. However, their model does not vary intrinsic fitness. If intrinsic fitness varies widely, a fitness-equalizing mechanism such as the JCH is necessary to impede competitive exclusion. Further unexplored complications are that JCEs vary in time and space (Janzen, 1972; Inman-Narahari *et al.*, 2016; LaManna *et al.*, 2016). While these factors require attention, it is unlikely that they would affect either JCE functional form more strongly than the other.

Results of this paper relate to empirically based criticisms of the JCH. Conspecific adults of common species in tropical forests persist close in space (less than 20 meters apart; (Hubbell, 1980)). Under the fixed neighborhood model, as Chisholm & Fung (2020) note, this spacing inhibits the feasible neighborhood effect size if JCEs are strong. However, for the additive model, strong JCEs fall to low values when seedlings are 20 meters from a single conspecific adult tree (Fig. 1B). Indeed, the additive predation model strongly facilitates coexistence precisely because the strength of JCEs increases when conspecifics are close in proximity (Fig. 1E) and outputs from the Spatially Explicit Model indicate that conspecific adults of common species are often within 20 meters of each other (Appendix D). Realistic dispersal kernels would also likely decrease the distance between conspecifics. Furthermore, habitat partitioning and JCEs can operate simultaneously to promote coexistence (Stump & Chesson, 2015). Conspecifics may persist in space closer than expected from a pure JCE model if a species’ fitness varies by habitat type and habitats exhibit spatial auto-correlation. Future studies should examine the interacting effects of auto-correlated habitat types, distant-dependent predation, and dispersal limitation.

It is also important to better characterize the nature of distance and density-dependent predation. For theoretical studies, this includes explicit modeling of seed/seedling natural enemies and their interactions (but see Schroeder *et al.* (2020)), examining the degree to which imperfect seed and seedling predator specialization affects diversity (but see Sedio & Ostling (2013)), and explicit modeling of multiple tree life history stages and their distance and density-dependent interactions. Empirical studies should aim to better quantify the key parameters that determine the strength of JCEs. The term that defines the strength of JCEs in this study, *a*, describes the sum of all mortality that may occur throughout the development of a non-adult tree via JCEs. Careful long-term measurements of inter-life history stage interactions are necessary to precisely evaluate this quantity – doing so is not a trivial task (Detto *et al.*, 2019). Quantification of species-specific values for the distance decay parameter (*ν*) and inter-specific fitness variation are also necessary to confirm or refute the efficacy of the JCH.

In conclusion, empirical evidence indicates that JCEs occur in a number of communities. I show, contrary to recent developments, the JCH is capable of preventing competitive exclusion under high levels of inter-specific fitness variation. The remaining question is whether the parameter space that corresponds to coexistence is realized in nature.

## Acknowledgments

This work was supported by the NSF NRT program, grant #DGE-1735359. I thank J. Timothy Wootton, Catherine A Pfister, Mercedes Pascual, Trevor Price, and Inés Moldavsky for insightful discussion, helpful comments, and feedback on the manuscript.

