## Appendix A for "The functional form of specialized predation dramatically affects whether Janzen-Connell effects can prevent competitive exclusion"

### Appendix A: Fixed neighborhood model and approximation

July 10, 2021

**4 Derivation of invasion criteria** **9**

**5 Figures** **14**

#### 1 Introduction

In this appendix, I derive and analyze the fixed neighborhood model from the main text. This appendix is composed of three sections: **(1)** I introduce a spatially explicit model (SEM) for the fixed neighborhood model. Then, I demonstrate that taking the expected seedling abundances on each patch yields the ODE model discussed in the main text. **(2)** To asses the accuracy of the ODE approximation, I provide outputs of the ODE model and the SEM model under the same parameterizations. I show the outputs are very similar, hence demonstrating that the ODE is a sufficiently accurate approximation of the SEM.**(3)** I provide the derivation of the invasion criteria approximation of the fixed neighborhood model (equation (8) in the main text).

#### 2 SEM and derivation of ODE approximation

In this section, I assume the reader is familiar with the model discussed in the main text. First, I discuss the SEM, briefly reviewing seed/seedling treatment from the main text. Then, I show the derivation of the ODE approximation.

##### 2.1 SEM

Using the similar methods of recent JCH-type models (Levi *et al.*, 2019; Chisholm & Fung, 2020) I developed a spatially explicit model. The model depicts a community with $M$ total patches modeled as an $L \times L$ grid ($L \times L = L^2 = M$). A torus was used to avoid edge effects. A single tree is present on every patch of the torus. At each time step, each tree has a probability $\delta$ of death and tree replacement occur via a lottery based on seedling abundances on each patch. As noted in the main text, seedling abundances are defined by

$$
\begin{aligned}
S_{i,i} &= Y_i\big[(1-d) + p_i d\big] J_{i,i}(p_i) \\
S_{i,k} &= Y_i p_i d J_{i,k}(p_i) \\
S_{all,i} &= \sum_{n=1}^{N} S_{n,i}
\end{aligned}
\tag{A.1}
$$

I consider when JCEs kill a fixed proportion of a species' seedlings on the nearest $E_n$ patches as defined by the Moore neighborhood. $E_n = 8$ is indicates a $3 \times 3$ Moore neighborhood, $E_n = 24$ indicates a $5 \times 5$ Moore neighborhood, and $E_n = 49$ indicates a $7 \times 7$ Moore neighborhood. Then, for the fixed neighborhood model,

$$J_{i,i}(p_i) = e^{-a_i}$$
$$J_{i,k}(p_k) = \begin{cases} 1, & \text{if } i \notin E_n \\ e^{-a_i}, & \text{if } i \in E_n \end{cases} \tag{A.2}$$

This yields the following seedling abundances :

$$S_{i,i} = Y_i\big[(1-d) + p_i d\big]e^{-a}$$
$$S_{i,k} = \begin{cases} Y_i d p_i, & \text{if } i \notin E_n \\ Y_i d p_i e^{-a}, & \text{if } i \in E_n \end{cases} \tag{A.3}$$
$$S_{all,i} = Y_i\big[(1-d) + dp_i\big]e^{-a} + \sum_{k \in E_n} Y_k p_k d e^{-a} + \sum_{j \notin E_n} Y_j p_j d$$

where $S_{x,y}$ is the seedling abundance of species $x$ on a patch occupied by species $y$. $i \in E_n$ is the condition that a conspecific tree of species $i$ is inside the neighborhood of a patch occupied by a heterospecific tree, $j$ and $i \notin E_n$ is when $i$ is not within the neighborhood. The lottery is determined by the relative seedling abundances. Let $P(x,y)$ be the probability species $x$ colonizes a patch previously occupied by species $y$. Then, $P(i,i) = S_{i,i}/S_{all,i}$ and $P(i,k) = S_{i,k}/S_{all,k}$. Patch-specific JCEs were determined by the closest $E_n$ neighbors on the torus (the Moore neighborhood).

##### 2.1.1 ODE Model

The goal of this section is to derive the approximate ODE

$$\frac{dp_i}{dt} = \delta\left[\frac{\mathbb{E}[S_{i,i}]}{\mathbb{E}[S_{all,i}]}p_i + \sum_{k\neq i}\frac{\mathbb{E}[S_{i,k}]}{\mathbb{E}[S_{all,k}]}p_k - p_i\right] \tag{A.4}$$

that captures the behavior of the SEM. The SEM neighborhood effect is implemented based on the nearest $E_n$ neighbors that, computationally, is stored in a matrix (the Moore neighborhood around a central point). A deterministic ODE model is by definition spatially implicit. Therefore, it is necessary to use an approximation of the terms that does not require spatial information. In particular, the quantity $S_{i,k}/S_{all,k}$, which contains the terms

$$S_{i,k} = \begin{cases} Y_i dp_i, & \text{if } i \notin E_n \\ Y_i dp_i e^{-a}, & \text{if } i \in E_n \end{cases} \tag{A.5}$$

and

$$S_{all,i} = \sum_{n=1}^{N} S_{n,i} \tag{A.6}$$

must be approximated as spatially implicit terms. There are several means of accomplishing this, but the simplest method is taking the expected expected seeds abundances on each patch. Assuming this is a good approximation (see the "Comparison between ODE model and SEM"), note that by the linearity of expectation,

$$\begin{aligned} \mathbb{E}[S_{all,i}] &= \mathbb{E}\left[\sum_{n=1}^{N} S_{n,i}\right] \\ &= \mathbb{E}\left[S_{i,i} + \sum_{n\neq i}^{N} S_{n,i}\right] \\ &= \mathbb{E}[S_{i,i}] + \sum_{n\neq i}\mathbb{E}\left[S_{n,i}\right] \end{aligned} \tag{A.7}$$

$\mathbb{E}[S_{i,i}]$ does not contain spatial information, as $\mathbb{E}[S_{i,i}] = \mathbb{E}[Y_i\big[(1-d)+p_i d\big]e^{-a}] = Y_i\big[(1-d)+p_i d\big]e^{-a}$. Therefore, it suffices to calculate the expected abundance of $S_{i,k}$, from which the other terms of interest (e.g. $S_{all,i}$) can be calculated. The expected seedling abundance on a given patch is equal to the number of seedlings on non-JCE affected patches multiplied the probability it does not experience JCEs added to the number of seedling on JCE affected patches multiplied the probability it does experience JCEs:

$$\mathbb{E}[S_{i,k}] = \mathrm{P}[i \notin E_n] \times S_{i,k}^{i \notin E_n} + \mathrm{P}[i \in E_n] \times S_{i,k}^{i \in E_n} \tag{A.8}$$

First, it is necessary to define $E_n$. As noted in the main text, if the neighborhood is of radius $r$ meters and assuming trees are of density $\gamma$ per square meter, there will be on average $\gamma\pi r^2$ individuals within the neighborhood defined by $r$. Therefore, $E_n = \gamma\pi r^2$.

The calculation of $\mathbb{E}[S_{i,k}]$ is as follows. Consider a single patch occupied by species $k$. It is necessary to calculate the probability an adult of species $i$ is within $E_n$. Recall that $\gamma$ is the density of trees in square meters, $r$ is the distance in meters from the local patch that defines the neighborhood effect, and $p_i$ be the proportion of species $i$ in the population. Assuming species are approximately randomly distributed, the probability that a conspecific is within the radius $r$ can be captured with a 2D Poisson process (i.e. a spatial Poisson process) with a rate parameter of $\lambda = p_i\gamma$. The waiting time of first event of a 2D Poisson process is exponentially distributed such that

$$\mathrm{P}[i \notin E_n] = e^{-\lambda A} \tag{A.9}$$

where $A$ is the area of the circle of radius $r$ stemming from the focal patch. Therefore $A = \pi r^2$.

Thus, $\lambda A = p_i \gamma \pi r^2$ and

$$\begin{aligned} \mathrm{P}[i \notin E_n] &= e^{-p_i \gamma \pi r^2} \\ \mathrm{P}[i \in E_n] &= 1 - e^{-p_i \gamma \pi r^2} \end{aligned} \tag{A.10}$$

Letting $\gamma \pi r^2 = E_n$ and using the above probabilities, the expected seedling value of $S_{i,k}$ is:

$$\begin{aligned} \mathbb{E}[S_{i,k}] &= \mathrm{P}[i \notin E_n] \times S_{i,k}^{i \notin E_n} + \mathrm{P}[i \in E_n] \times S_{i,k}^{i \in E_n} \\ &= Y_i d p_i \big[ e^{-p_i E_n} + e^{-a}(1 - e^{-p_i E_n}) \big] \end{aligned} \tag{A.11}$$

Plugging this value into the post-JCE seedling abundance terms yields

$$\begin{aligned} S_{i,i} &= Y_i \big[ (1-d) + d p_i \big] e^{-a} \\ S_{i,k} &= Y_i d p_i \big[ e^{-p_i E_n} + e^{-a}(1 - e^{-p_i E_n}) \big] \end{aligned} \tag{A.12}$$

also noting that $S_{all,i} = \sum_{n=1}^{N} S_{n,i}$. This is identical to equation (5) from the main text.

#### 3 Comparison between ODE model and SEM

In this section, I describe simulations that compare the ODE model to the SEM. I demonstrate that the SEM and ODE model, to a first approximation, yield the same outputs of species abundance, species richness, and effective species richness. I compare the ODE and SEM outputs of the fixed neighborhood model under the relevant scenario examined in the main text: interspecific variation in intrinsic fitness ($Y$). I provide 27 comparisons of the SEM to the ODE (9 cases in which $E_n = 8$, 9 cases in which $E_n = 24$, and 9 cases in which $E_n = 49$). Parameter values were chosen to encapsulate the range of parameters shown in the main text (see Fig. A1).

##### 3.1 ODE modification for SEM comparison

The SEM takes place on a grid (a $275 \times 275$ patches modeled on a torus) on which each patch is occupied by a single adult tree. Therefore, there is a fixed distance between individuals such that species are distributed regularly (with a given fixed distance). In contrast, the ODE described above assumes that species are distributed randomly in continuous space such that $E_n$ ($E_n = \gamma\pi r^2$) can be any real number. However, for the SEM, it is necessary to the number of neighbors as a discrete number. Therefore, I make the below minor modification to the ODE.

Recall that the expected effect of specialized predation in the fixed neighborhood model is

$$e^{-p_i E_n} + e^{-a}(1 - e^{-p_i E_n})$$

To transform this into term compatible with the ODE, note that $e^{-piE_n} \approx (1 - p_i)^{E_n}$. This logical – this probability that species $k$ falls into the neighborhood effect area is mathematically equivalent to $E_n$ Bernoulli trials. To see this, let $x = (1-p_k)^{E_n}$. Taking the log of each side yields $\log[1 - x] = E_n \log[(1 - p_k)]$. Noting that $\log[1 - m] \approx -m$ for small $m$ and assuming $p_k$ is fairly small yields the expression $\log[x] \approx -E_n p_k$. Taking the exponent of each side yields $x \approx e^{-p_k E_n}$, which is what we wanted. For the SEM, which directly considers a discrete neighborhood of $E_n$ patches adjacent to focal patch, the Bernoulli trial approximation is more appropriate than the exponential terms (albeit almost quantitatively identical). Using this yields

$$e^{-p_i E_n} + e^{-a}(1 - e^{-p_i E_n}) \approx (1 - p_i)^{E_n} + e^{-a}\big(1 - (1 - p_i)^{E_n}\big) \tag{A.13}$$

I use this for the ODE-SEM comparison.

#### 3.2 ODE and SEM parameterization

Each SEM simulation began with 300 species at equal abundance, with individuals randomly distributed throughout the community. Simulations were conducted on a $275 \times 275$ torus (thus containing $275^2$ individual trees). I use the following parameters: $Y \sim$ lognormal$[\mu = 0, \sigma_Y]$ with $\sigma_Y \sim \{0.1, 0.45, 0.8\}$ and $a \sim \{1, 2.75, 4.5\}$. In all simulations, $d = 1$. I tested each of the

9 parameter combinations with $E_n = 8$, 24, and 49. This generated 27 outputs. Simulations were run for about 65 generations, sufficient time for the community to approximately reach equilibrium without drift dominating the dynamics of the lower abundance species. However, it was noticed that the cases in which $\sigma_Y = 0.1$ had longer transient times. These were run for 125 generation instead. See Fig. A5-A7 for examples of the transient dynamics. I also ran a set of 27 ODE simulation using the same parameterizations as the SEM. I compared the outputs of the SEM and ODE model in terms of species diversity, effective species richness (the exponential of Shannon Diversity), and abundance. I considered a species to be extinct if it had less than 5 individuals at any point of the simulation. This was implemented directly in the SEM; for the ODE model, I assumed a species, $i$, to be extinct if $p_i^* < 5/275^2$ where $p_i^*$ is the equilibrium proportion of species $i$. This cutoff was put into palace to reduce the probability of counting a species from the SEM with a negative growth rate near extinction as persisting. Note that these simulations do not attempt to demonstrate the long-term resistance against extinction due to drift. Rather, they demonstrate that the ODE model and SEM yield similar outputs in species expected abundance and total diversity given the same parameterization.

#### 3.3 Results of comparison

As can be seen in Fig. A2, the ODE and SEM yield very similar results in species richness and effective species richness. To quantify the quality of the approximation, I used the mean percentage error (MPE) as calculated by

$$\text{MPE} = \frac{1}{S}\sum_{i=1}^{S}\left|\frac{D_{\text{SEM},i} - D_{\text{ODE},i}}{D_{\text{SEM},i}}\right| \times 100 \tag{A.14}$$

where $S$ is the total number of simulations, $D_{\text{SEM},i}$ is the diversity of the $i_{th}$ SEM simulation, and $D_{\text{ODE},i}$ is the diversity the $i_{th}$ ODE model simulation. Over all simulations, MPE = 8.70% for species richness and 9.34% for effective species richness. Overall, the ODE provides a highly similar, albeit non-exact, estimation of species diversity. Any errors in diversity are trivial relative to the difference between the outputs of JCE functional forms. The SEM and ODE model also produced highly similar species abundances (see Fig. A3-A4).

#### 4 Derivation of invasion criteria

In this section, I derive the approximate invasion criteria of the fixed neighborhood effect model when species experience inter-specific variation in intrinsic ($Y$) and $d = 1$. The invasion criteria of an invader can be expressed as when the per capita growth rate as $p_i \to 0$. Letting

$$J_k(p_k) = e^{-p_k E_n} + e^{-a}(1 - e^{-p_k E_n}) \tag{A.15}$$

The per capita growth rate of species $i$, substituting in the seedling abundance values, is

$$\begin{aligned} \frac{1}{p_i}\frac{dp_i}{dt} = r_i = \delta\Bigg[ & \frac{Y_i\big[(1-d)+p_i d\big]}{Y_i\big[(1-d)+p_i d\big] + \sum_{k\neq i} Y_k p_k d J_k(p_k)} \\ & + Y_i d \sum_{m\neq i} \frac{1}{Y_m e^{-a}\big[(1-d)+p_m d\big] + \sum_{k\neq m} Y_k p_k d J_k(p_k)} p_m - 1\Bigg] \end{aligned} \tag{A.16}$$

Species $i$ can invade is this quantity if positive when it is rare ($p_i \to 0$). When $d = 1$ (the case of interest), the above reduces to

$$Y_i \sum_{m\neq i} \frac{p_m}{Y_m e^{-a} p_m + \sum_{k\neq m} Y_k p_k J_k(p_k)} > 1 \tag{A.17}$$

To simplify the above equation, I ignore the term $Y_m e^{-a} p_m$ in the denominator and incorporate an additional term the summation, yielding:

$$Y_i \sum_{m\neq i} \frac{p_m}{\sum_{k\neq i} Y_k p_k J_k(p_k)} > 1 \tag{A.18}$$

This simplification is equivalent to making the species identity of the tree previously occupying a patch (the tree that dies) irrelevant (i.e., JCEs only result from trees nearby the patch rather than the previous occupant of the patch). As long as neighborhood effects are somewhat strong – that is, so long as $E_n$ is not very small – this assumption should not meaningfully affect the invasion criteria. Numerical simulations show that for the values considered in the main text this simplification does not qualitatively affect results (see Appendix C).

Importantly, the denominator of equation (A.18) is no longer directly dependent on $m$. That is, equation (A.18) can be rewritten as

$$Y_i\left(\sum_{m\neq i} p_m\right)\left(\frac{1}{\sum_{k\neq i} Y_k p_k J_k(p_k)}\right) > 1 \tag{A.19}$$

Because, by definition, $\sum_{m\neq i} p_m = 1$, the above equation can be rewritten as

$$Y_i > \sum_{k\neq i} Y_k p_k J_k(p_k) \tag{A.20}$$

Substituting the appropriate value for $J_k(p_k)$ and letting $C = 1 - e^{-a}$, the invasion criteria becomes

$$Y_i > \sum_{k\neq i} Y_k p_k \big(1 - C(1 - e^{-p_k E_n})\big) \tag{A.21}$$

Taking the 1st order Taylor expansion of $p_k\big(1-C(1-e^{-p_k E_n})\big)$ around the point $1/N$ where $N$ is the number of species in the resident community yields a close approximation of the expression so long as variation in $p_k$ is not very large. Thus,

$$p_k\big(1 - C(1 - e^{-p_k E_n})\big) \approx \frac{Ce^{-\frac{E_n}{N}} E_n}{N^2} + p_k\left[1 - C\left(1 - e^{-\frac{E_n}{N}}\left(1 - \frac{E_n}{N}\right)\right)\right] \tag{A.22}$$

The summation can be broken up into the two additive terms:

$$\sum_{k\neq i} Y_k \frac{Ce^{-\frac{E_n}{N}}E_n}{N^2} + \sum_{k\neq i} Y_k p_k \left[1 - C\left(1 - e^{-\frac{E_n}{N}}\left(1 - \frac{E_n}{N}\right)\right)\right] \tag{A.23}$$

The first term is trivial to calculate:

$$\begin{aligned}\sum_{k\neq i} Y_k \frac{Ce^{-\frac{E_n}{N}}E_n}{N^2} &= \frac{Ce^{-\frac{E_n}{N}}E_n}{N}\frac{1}{N}\sum_{k\neq i} Y_k \\ &= \overline{Y}Ce^{-\frac{E_n}{N}}\frac{E_n}{N}\end{aligned} \tag{A.24}$$

where $\overline{Y}$ is the mean intrinsic fitness of the resident community.

The second term in the summation can be expressed by noting the property

$$\frac{1}{N}\sum_{m=1}^{N} A_m B_m = \overline{A} \times \overline{B} + \mathrm{Cov}(A, B) \tag{A.25}$$

Using this property and substituting $A$ and $B$ with $p$ and $Y$ yields

$$\begin{aligned}\sum_{k\neq i} Y_k p_k \left[1 - C\left(1 - e^{-\frac{E_n}{N}}\left(1 - \frac{E_n}{N}\right)\right)\right] &= \left[1 - C\left(1 - e^{-\frac{E_n}{N}}\left(1 - \frac{E_n}{N}\right)\right)\right] N \frac{1}{N}\sum_{k\neq i} Y_k p_k \\ &= \left[1 - C\left(1 - e^{-\frac{E_n}{N}}\left(1 - \frac{E_n}{N}\right)\right)\right] N \left(\frac{\overline{Y}}{N} + \mathrm{Cov}(p, Y)\right) \\ &= \left[1 - C\left(1 - e^{-\frac{E_n}{N}}\left(1 - \frac{E_n}{N}\right)\right)\right] \left(\overline{Y} + N\mathrm{Cov}(p, Y)\right)\end{aligned} \tag{A.26}$$

Just considering the $\overline{Y}$ term,

$$\overline{Y}\left[1 - C\left(1 - e^{-\frac{E_n}{N}}\left(1 - \frac{E_n}{N}\right)\right)\right] = \overline{Y}\left(1 - C\left(1 - e^{\frac{-E_n}{N}}\right)\right) - \overline{Y}Ce^{-\frac{E_n}{N}}\frac{E_n}{N} \tag{A.27}$$

Adding the $\overline{Y}$ terms (equations (A.24) and (A.27)) is

$$\begin{aligned}
\overline{Y}Ce^{-\frac{E_n}{N}}\frac{E_n}{N} + \overline{Y}\left(1 - C\big(1 - e^{\frac{-E_n}{N}}\big)\right) - \overline{Y}Ce^{-\frac{E_n}{N}}\frac{E_n}{N} &= \overline{Y}\left(1 - C\big(1 - e^{\frac{-E_n}{N}}\big)\right) \\
&\approx \overline{Y}\left(1 - C\frac{E_n}{N}\right) \\
&\approx \overline{Y}e^{-C\frac{E_n}{N}}
\end{aligned} \tag{A.28}$$

where the approximations come from the property noted in the above section that $e^{-x} \approx 1 - x$ if $x$ is somewhat small.

For the covariance term, the outside constant can be arranged as following (again recalling that $e^{-x} \approx 1 - x$ if $x$ is somewhat small):

$$\begin{aligned}
1 - C\left(1 - e^{-\frac{E_n}{N}}\left(1 - \frac{E_n}{N}\right)\right) &= 1 - C\big(1 - e^{-\frac{E_n}{N}}\big) - Ce^{-\frac{E_n}{N}}\frac{E_n}{N} \\
&\approx 1 - C\frac{E_n}{N} - C\frac{E_n}{N}\left(1 - \frac{E_n}{N}\right) \\
&= 1 - C\frac{E_n}{N}\left(2 - \frac{E_n}{N}\right) \\
&\approx e^{-C\frac{E_n}{N}\left(2 - \frac{E_n}{N}\right)} \\
&= e^{-C\frac{E_n}{N} - C\frac{E_n}{N} + C\frac{E_n^2}{N^2}} \\
&= e^{-C\frac{E_n}{N}}e^{-C\frac{E_n}{N} + C\frac{E_n^2}{N^2}}
\end{aligned} \tag{A.29}$$

Therefore, the invasion criteria can be written as

$$Y_i > \overline{Y}e^{-C\frac{E_n}{N}} + e^{-C\frac{E_n}{N}}e^{-C\frac{E_n}{N} + C\frac{E_n^2}{N^2}}N\text{Cov}(p, Y) \tag{A.30}$$

which can be rewritten as

$$Y_i > \overline{Y}e^{-\left(1 - e^{-a}\right)\frac{E_n}{N}}\left[1 + e^{-\left(1 - e^{-a}\right)\frac{E_n}{N} + \left(1 - e^{-a}\right)\frac{E_n^2}{N^2}}\frac{\text{Cov}(p, Y)}{\overline{Y}/N}\right] \tag{A.31}$$

I then make two simplifications. Firstly, because it is assumed that $E_n << N$ , then $(E_n/N)^2 \approx 0$. Secondly, once again note that $e^{-x} \approx 1 - x$ if $x$ is somewhat small. Therefore, $e^{-\left(1-e^{-a}\right)\frac{E_n}{N}} \approx 1 - \left(1 - e^{-a}\right)\frac{E_n}{N}$. Therefore, the above can be rewritten as

$$Y_i > \underbrace{\overline{Y}}_{\substack{\text{mean}\\ \text{fitness}}} \underbrace{e^{-\left(1-e^{-a}\right)\frac{E_n}{N}}}_{\text{mean JCE}} \left[1 + \underbrace{\frac{\text{Cov}(p, Y)}{\overline{Y}/N}}_{\substack{\text{covariance}\\ \text{term}}} \underbrace{\left(1 - \left(1 - e^{-a}\right)\frac{E_n}{N}\right)}_{\substack{\text{covariance}\\ \text{coefficient}}}\right] \tag{A.32}$$

which is the same as equation (8) from the main text.

### 5 Figures

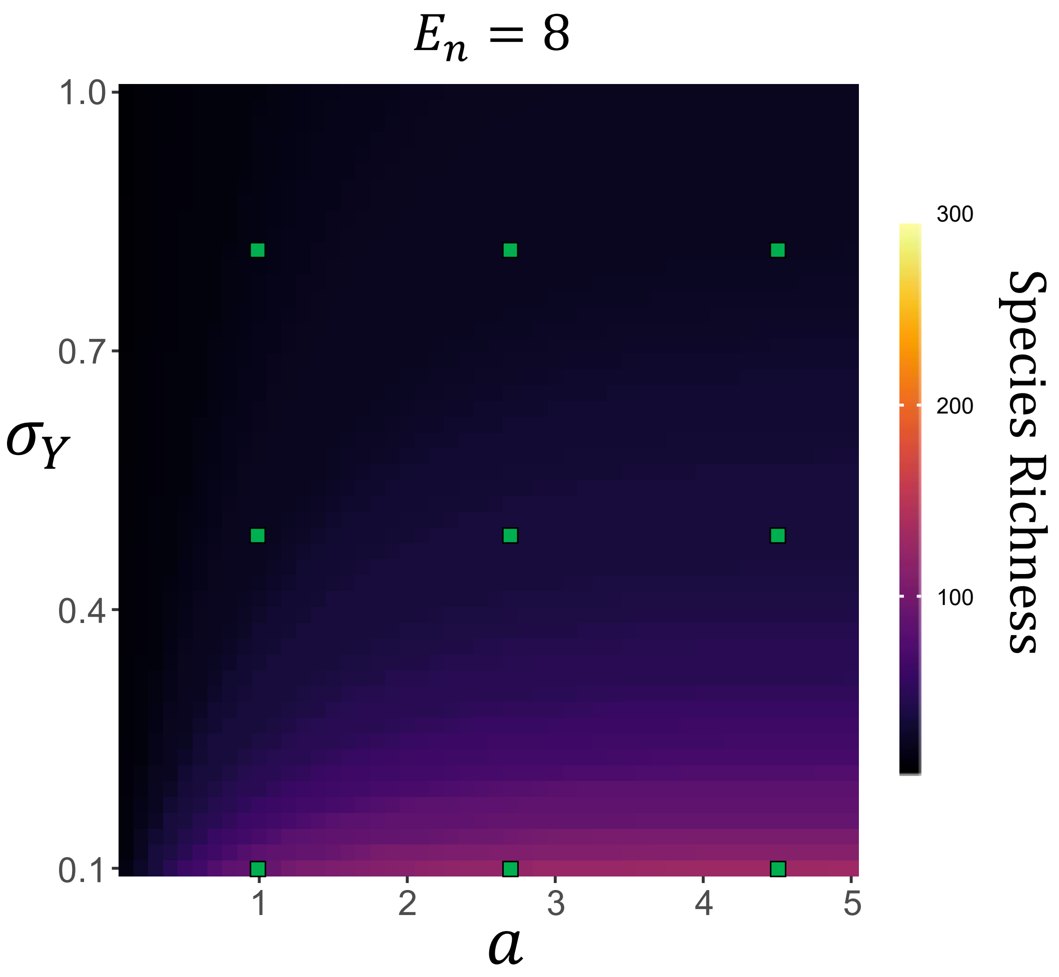

**Fig. A1:** Visualization of the parameter values used for the SEM. Green squares depict the parameter values run in each SEM simulation. Values are superimposed on the heatmaps from the main text to show the extent of parameter space examined. $E_n = 8$ is noted for consistency with the figure from the main text, but the same parameters were tested for the $E_n = 24$ and $E_n = 49$ cases as well.

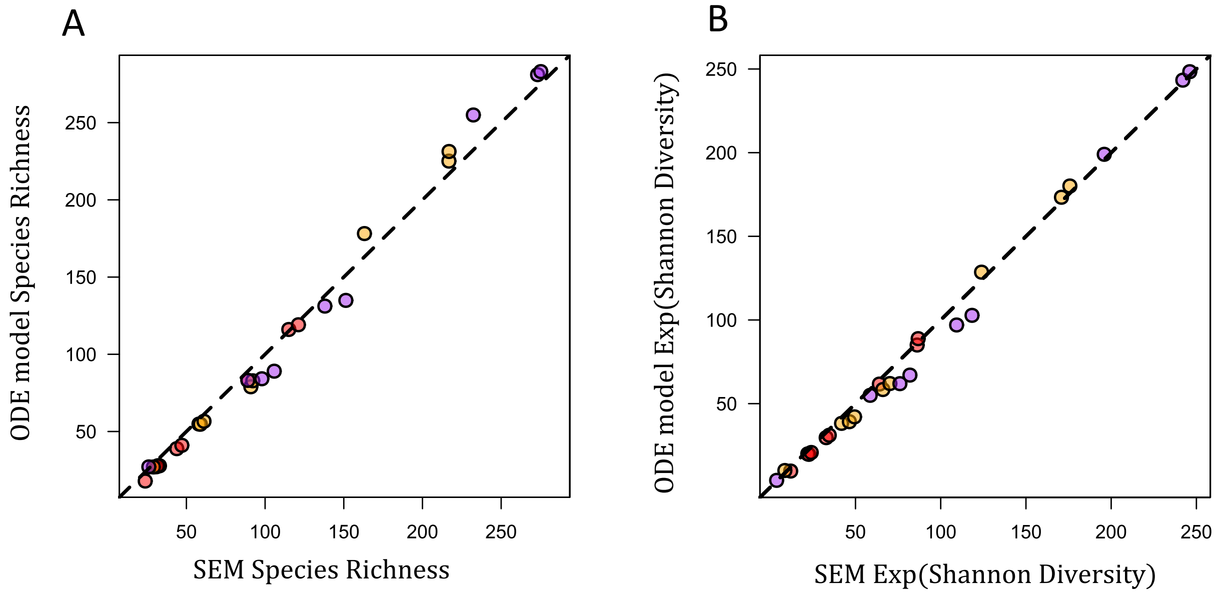

**Fig. A2:** ODE model validation. The figures compare species richness and effective species richness (the exponential of Shannon Diversity) between SEM and ODE model simulations under identical parameterizations. The dashed line is the one-to-one line (points on the line represent when the SEM and ODE yield the exact same diversity output). Red points are when $E_n = 8$, orange/yellow points are when $E_n = 24$, and purple points are when $E_n = 49$. To a first approximation, the ODE model yields the same output as the SEM in both diversity metrics.

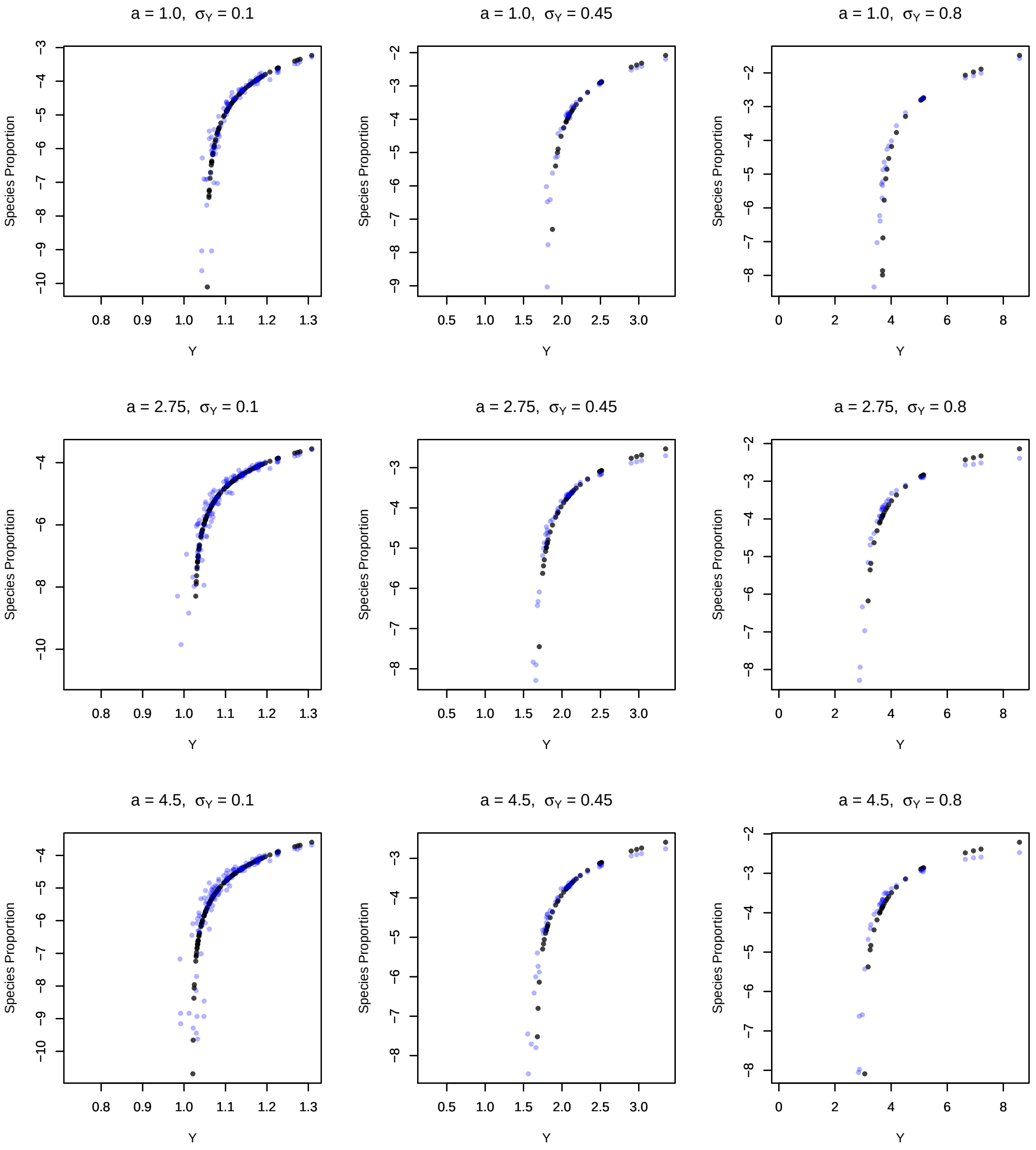

**Fig. A3:** Comparisons between identical parameterizations of the ODE approximation (black) and SEM (blue) outputs under nine parameter values when species vary in intrinsic fitness ($Y$). The$y$-axis depicts the log-proportion of each species and the $x$-axis depicts $Y$ of each species. These parameter values span the most of the parameter space explored in Fig. 4 of the main text. In all plots, $E_n = 8$, and $d = 1.0$. Other relevant parameters are listed on each plot.

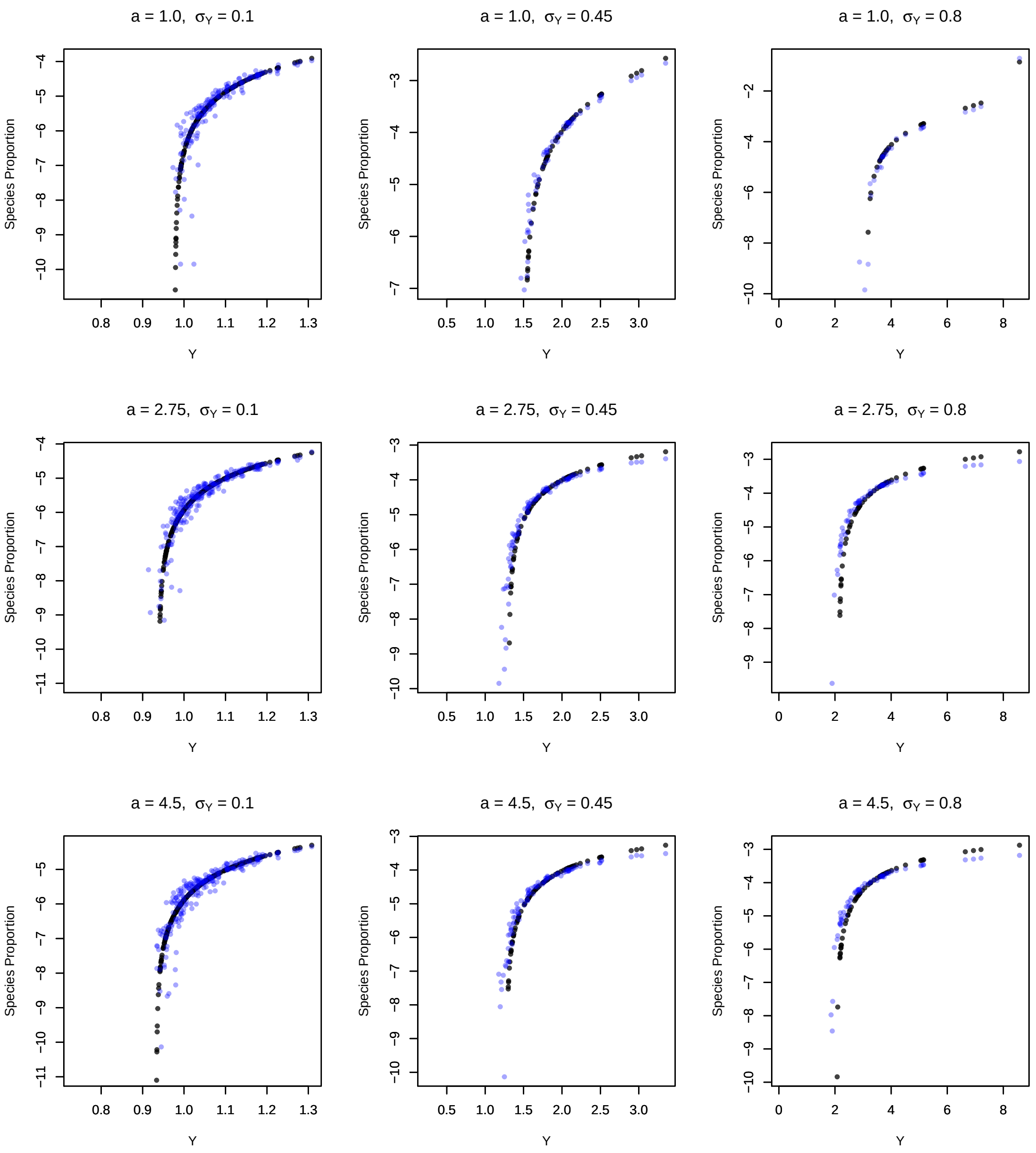

**Fig. A4:** The same format as Fig. A3, but with $E_n = 24$.

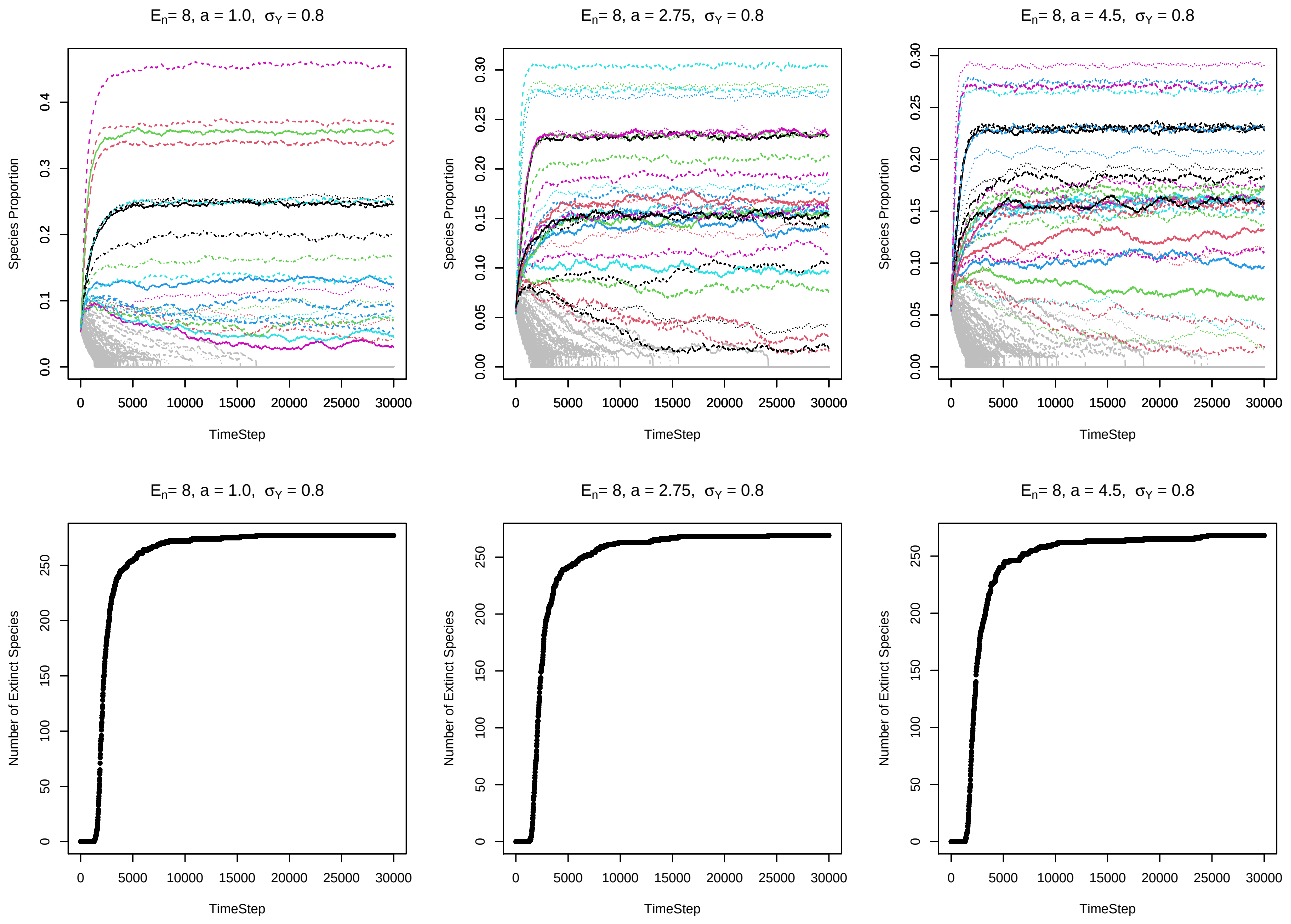

**Fig. A5:** Examples of the SEM simulation time series outputs. The top row shows examples of the time series outputs of the SEMs. The $x$-axis is time and the $y$-axis is each species' proportion. Proportions have been square-root transformed to aid visualization. Colored trajectories indicate species that persisted throughout the simulation; grey trajectories indicate species that went extinct. In these examples, $Y$ varies between species. Parameters are listed on each plot. Dynamics as shown are typical examples from the SEMs. Most species settle into a relatively stable pseudo-equilibrium, while lower abundance species fluctuate due to drift. The bottom row shows the number of extinct species in the community as a function of time. Each panel corresponds to the plot above it. As can be seen, most species that go extinct do so in the early stages of the dynamics. Therefore, the vast majority of persisting species likely persist deterministically. $E_n = 8$ in a simulations.

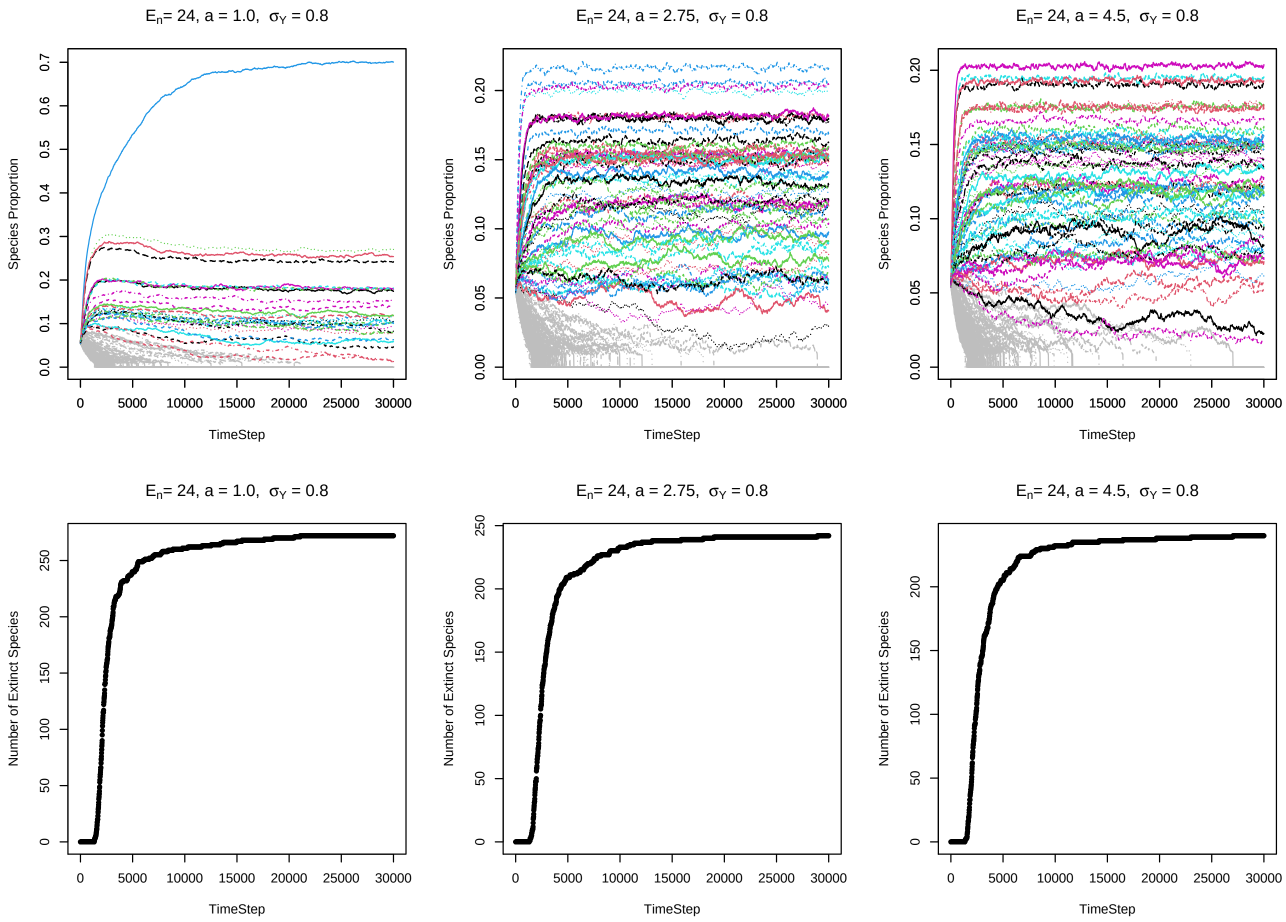

**Fig. A6:** The same format as Fig. A5, but with $E_n = 24$.

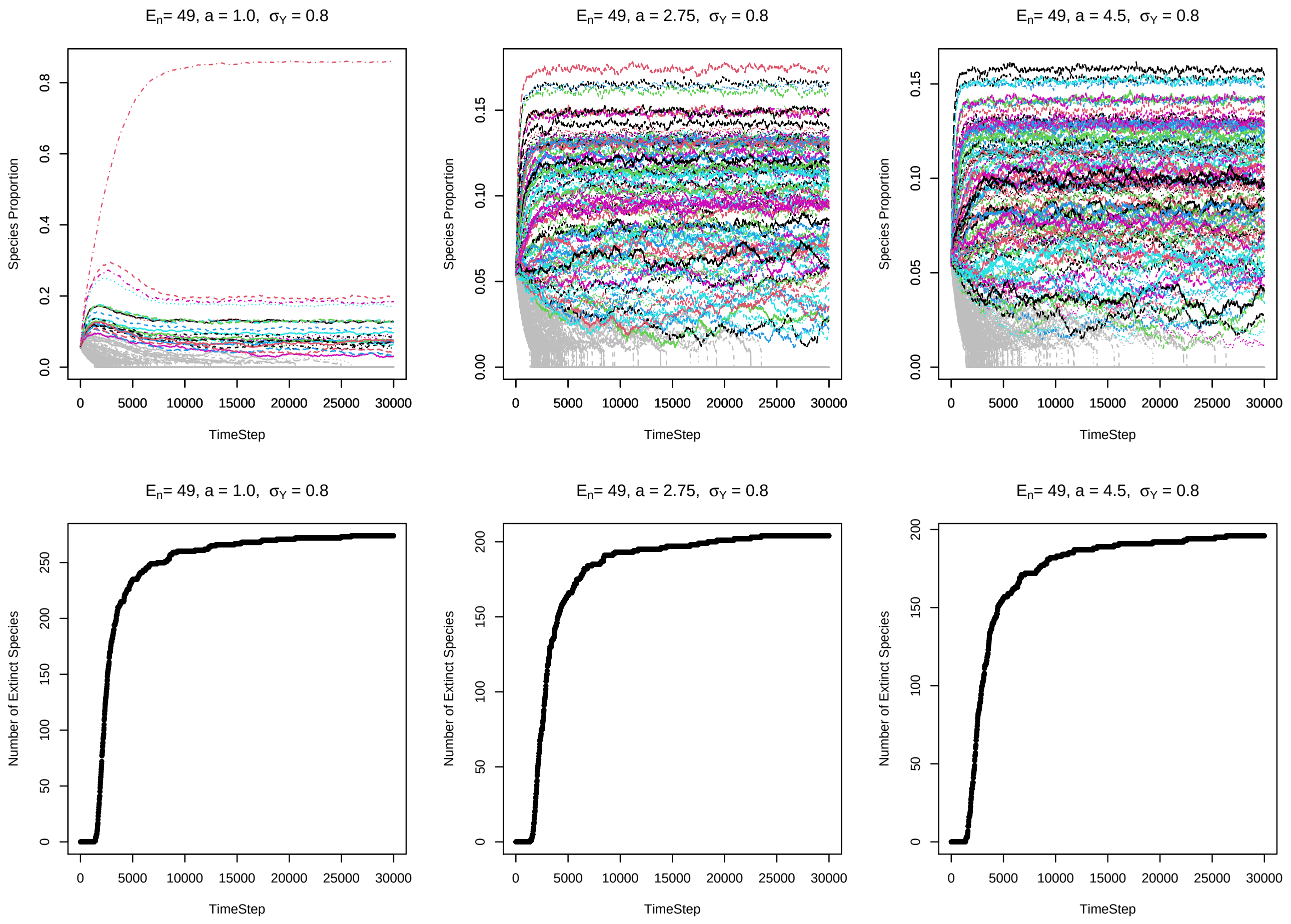

**Fig. A7:** The same format as Fig. A5, but with $E_n = 49$.
