## Appendix B for "The functional form of specialized predation dramatically affects whether Janzen-Connell effects can prevent competitive exclusion"

### Appendix B: additive predation model SEM and ODE approximation

July 10, 2021

**4 Derivation of invasion criteria** **9**

**5 Figures** **13**

### 1 Introduction

In this Appendix, I analyze the additive predation model presented in the main text. This appendix is composed of three main sections: **(1)** I introduce a spatially explicit model (SEM) of the additive predation model. I then demonstrate that taking the expected seedling abundances on each patch yields the ODE model discussed in the main text. **(2)** I provide outputs of the ODE model and the SEM model under the same parameterizations. I show the outputs are very similar, hence demonstrating that the ODE is a sufficiently accurate approximation of the SEM. **(3)** I provide the derivation of the invasion criteria approximation of the additive predation model (equation (7) in the main text). The structure of this appendix is identical to that of Appendix A.

### 2 SEM and derivation of ODE approximation

In this section, I discuss the SEM, briefly reviewing seed/seedling treatment from the main text and show the derivation of the ODE approximation. I assume the reader is familiar with the general model discussed in the main text.

#### 2.1 SEM

I developed a spatially explicit model that integrates the additive predation model. The model consists of a community on a grid of $L \times L$ patches ($M$ total patches, $M = L^2$) modeled as a torus to avoid edge effects. A single tree is present on every space on the grid. At each time step, each tree dies with probability $\delta$ and tree replacement occur via a lottery based on seedling abundances on each patch. I assume predation pressure increases linearly as a function of local conspecific density and that the predation effect of conspecific proximity decreases exponentially with distance. As noted in the main text, seedling abundances are defined by

$$
\begin{aligned}
S_{i,i} &= Y_i\big[(1-d) + N_i d\big] J_{i,i}(N_i) \\
S_{i,k} &= Y_i N_i d J_{i,k}(N_i) \\
S_{all,i} &= \sum_{n=1}^{N} S_{n,i}
\end{aligned}
\tag{B.1}
$$

where, for the additive predation case,

$$
\begin{aligned}
J_{i,i}(p_i) &= \exp\Bigg[-a_i\Bigg(1 + \sum_{x_i=1}^{Mp_i} e^{-x_i/\nu}\Bigg)\Bigg] \\
J_{i,k}(p_i) &= \exp\Bigg[-a_i \sum_{x_i=1}^{Mp_i} e^{-x_i/\nu}\Bigg]
\end{aligned}
\tag{B.2}
$$

where $x_i$ is the distance of the $x_{th}$ closest individual of species $i$ to the focal patch. Seedling abundances are then determined by the following equations:

$$
\begin{aligned}
S_{i,i} &= Y_i\big[(1-d) + dp_i\big] \exp\Bigg[-a\Bigg(1 + \sum_{x_i=1}^{Mp_i} e^{-x_i/\nu}\Bigg)\Bigg] \\
S_{i,k} &= Y_i p_i d \exp\Bigg[-a_i \sum_{x_i=1}^{Mp_i} e^{-x_i/\nu}\Bigg] \\
S_{i,all} &= Y_i\big[(1-d) + dp_i\big] \exp\Bigg[-a\Bigg(1 + \sum_{x_i=1}^{Mp_i} e^{-x_i/\nu}\Bigg)\Bigg] + \sum_{k\neq i} Y_k p_k d \exp\Bigg[-a_i \sum_{x_i=1}^{Mp_i} e^{-x_i/\nu}\Bigg]
\end{aligned}
\tag{B.3}
$$

where $S_{X,Y}$ is the seedling abundance of species $X$ on a patch occupied by species $Y$. $\nu$ defines rate at which predataion declines with distance (higher $\nu$ indicates a lower rate of predation decay). The lottery is determined by the relative seedling abundances. Let $P(X, Y)$ be the probability species $X$ colonizes a patch previously occupied by species $Y$. Then, $P(i,i) = S_{i,i}/S_{all,i}$ and $P(i,k) = S_{i,k}/S_{all,k}$. For the SEM, patch-specific JCEs are determined by the euclidean distances of species surrounding each site. Distances between patches are measured by the distance between the center point of each patch.

#### 2.2 ODE Model

The goal of this section is to derive the approximate ODE

$$\frac{dp_i}{dt} = \delta \left[ \frac{\mathbb{E}[S_{i,i}]}{\mathbb{E}[S_{all,i}]} p_i + \sum_{k \neq i} \frac{\mathbb{E}[S_{i,k}]}{\mathbb{E}[S_{all,k}]} p_k - p_i \right] \tag{B.4}$$

that captures the behavior of the SEM. The SEM additive predation function is implemented based on the proximity of neighboring species to a patch that, computationally, is stored in a matrix. A deterministic ODE model is by definition spatially implicit. Therefore, it is necessary to use an approximation of the terms that do not require spatial information. In particular, the quantity $S_{i,k}/S_{all,k}$, which contains the term

$$a_i \sum_{x_{i,k} \in M} e^{-x_{i,k}/\nu} \tag{B.5}$$

henceforth the "predation rate". This quantity must be approximated as a spatially implicit term. To do so, I simply calculate the expected seedling predation pressure each species experiences.

Consider a single patch occupied by species $k$. From that patch, individuals of species $i$ persists in space with a distribution of distances from the focal patch. Let $\gamma$ be the density of trees in square meters, let $x_m$ be the distance in meters of the $m_{th}$ closest individual of species $i$ from the focal patch, and let $p_i$ be the proportion of species $i$ in the population. Assuming species are approximately randomly distributed, the number of conspecifics of species $i$ within a given distance, $x$, from the focal patch can be captured with a 2D Poisson process (also known as a spatial Poisson process). The expected predation rate of species $i$ on a patch occupied by species $k$ is equal to $a_i \times 2\gamma\pi p_i \nu^2$. The proof is shown below – this proof is somewhat trivial (and longer than necessary) but one step is necessary for the subsequent section.

Let $x_m$ be the $m_{th}$ distance away from a focal patch occupied by species $k$. By the definition of a 2D Poisson process, the expected abundance of individuals of species $i$ between the $m_{th}$ and the $(m-1)_{th}$ distances is proportional to the area therein, $p_i\gamma\pi(x_m^2$ - $x_m^2)$. The expected predation pressure can be calculated as the sum of the expected number of conspecifics of species $i$ for each interval of $x_{m-1}$ to $x_m$ scaled by exponential decay with distance over all distances, or

$$a\gamma\pi p_i \sum_{m=1}^{\infty} \big(x_m^2 - x_{m-1}^2\big) e^{-x_m/\nu} \tag{B.6}$$

Let $m\Delta x = x_m$ where $\Delta x = x_m - x_{m-1}$. Then, the above summation can be written as

$$a\gamma\pi p_i \sum_{m=1}^{\infty} \Big( \big(m\Delta x\big)^2 - \big((m-1)\Delta x\big)^2 \Big) e^{-m\Delta x/\nu} \tag{B.7}$$

As $\Delta x$ becomes very small (meaning that space is considered continuously rather than discretely)

$$\begin{aligned} \lim_{\Delta x \to 0} \left[ a_i\gamma\pi p_i \sum_{m=1}^{\infty} \Big( \big(m\Delta x\big)^2 - \big((m-1)\Delta x\big)^2 \Big) e^{-m\Delta x/\nu} \right] &\to a_i\gamma\pi p_i \int_0^{\infty} \left( \frac{d}{dx} x^2 \right) e^{-x/\nu} dx \\ &= a_i \times 2\gamma\pi p_i \int_0^{\infty} x e^{-x/\nu} dx \\ &= a_i \times 2\gamma\pi p_i \nu^2 \end{aligned} \tag{B.8}$$

which is what we wanted. Therefore,

$$\mathbb{E} \left[ a_i \sum_{x_{i,k} \in M} e^{-x_{i,k}/\nu} \right] = a_i \times 2\gamma\pi p_i \nu^2 \tag{B.9}$$

Letting $E_A = 2\gamma\pi\nu^2$ and substituting this into the seedling abundance (equation (B.3)) yields

$$\begin{aligned} S_{i,i} &= Y_i \big[(1-d) + dp_i\big] e^{-a_i(1+p_iE_A)} \\ S_{i,k} &= Y_i dp_i e^{-a_i p_i E_A} \end{aligned} \tag{B.10}$$

where $S_{all,i} = \sum_{n=1}^{N} S_{n,i}$. This is the same as equation (6) in the main text.

#### 3 Comparison between ODE model and SEM model

In this section, I describe simulations that compare the ODE model to the SEM. I demonstrate that the SEM and ODE model, to a first approximation, yield the same outputs of species abundance and species richness. I provide 27 comparisons of the SEM to the ODE (9 cases in which $\nu = 5$, 9 cases in which $\nu = 7.5$, and 9 cases in which $\nu = 10$. Parameter values were chosen to encapsulate the range of parameters shown in the main text (see Fig. B1).

##### 3.1 ODE modification for SEM comparison

First, it is necessary to slightly modify the ODE approximation to match the structure of the SEM. From the above section, recall that the predation pressure term is calculated as

$$a\gamma\pi p_i \sum_{m=1}^{\infty} \left( \left(m\Delta x\right)^2 - \left((m-1)\Delta x\right)^2 \right) e^{-m\Delta x/\nu} \tag{B.11}$$

for which the limit of this above expressions is taken as $\Delta x \to 0$. This is logical when treating space as a continuous variable with respect to species location. However, in the SEM, each patch is a fixed distance apart because the community is modeled on a grid (torus) with $L \times L$ patches (and hence, $L \times L$ individuals). The fixed distance between patches is $\Delta x$ meters. Thus, to directly compare the ODE and SEM, the above equation must be evaluated for a value of $\Delta x$ that reflects the imposed spacing of the SEM. The distance between patches is simply the average distance between particles in 2 dimensions. This is a well-known relationship and can be easily calculated from $\gamma$, the tree density (in units individual per square meter). $\gamma$ is ratio of total individuals ($L^2$) in the community divided by the area of the community in meters. The total area of the community is $(\Delta x L)^2$ where $\Delta x$ is the distance between patches in meters. Thus

$L^2/(\Delta x L)^2 = \gamma$. Solving for $\Delta x$ yields

$$\Delta x = \frac{1}{\sqrt{\gamma}} \tag{B.12}$$

Substituting this value into the limit yields

$$\lim_{\Delta x \to \frac{1}{\sqrt{\gamma}}} \left[ a\gamma\pi p_i \sum_{m=1}^{\infty} \left( \left(m\Delta x\right)^2 - \left((m-1)\Delta x\right)^2 \right) e^{-m\Delta x/\nu} \right] \to a\pi p_i \frac{1+\exp(1+\frac{1}{\nu\sqrt{\gamma}})}{\left[1-\exp(1+\frac{1}{\nu\sqrt{\gamma}})\right]^2} \tag{B.13}$$

This yields a minor yet but quantifiable change to the ODE output. I use this value to compare the SEM and ODE.

#### 3.2 ODE and SEM parameterization

Each SEM simulation began with 300 species at equal abundance, with individuals randomly distributed throughout the community. Simulations were conducted on a 275 × 275 torus (thus containing $275^2$ individual trees). I used the following parameters: $Y \sim \text{lognormal}[\mu = 0, \sigma_Y]$ with $\sigma_Y \sim \{0.1, 0.45, 0.8\}$ and $a \sim \{1, 2.75, 4.5\}$. In all simulations, $\gamma = 0.20$ and $d = 1$. I tested each of the 9 parameter combinations with $\nu \sim \{5, 7.5, 10\}$. This generated 27 outputs. See Fig. B1 for a visualization of the parameter space explored. Simulations were run for about 65 generations, sufficient time for the community to approximately reach equilibrium without drift dominating the dynamics of the lower abundance species. See Figs. B6-B8 for typical outputs of the SEM time series dynamics. A corresponding set of 27 ODE simulations were run using the same parameterizations as the SEM. I compared the outputs of the SEM and ODE model in terms of species diversity, effective species richness (the exponential of Shannon Diversity), and abundance. I considered a species to be extinct if it had less than 5 individuals. This was implemented directly in the SEM; for the ODE model, I assumed a species, $i$, to be extinct if $p_i^* < 5/275^2$ where $p_i^*$ is the equilibrium proportion of species $i$. This cutoff was put into palace to reduce the probability of counting a species from the SEM with a negative growth rate nearing extinction as persisting.

##### 3.3 Results of comparison

The ODE and SEM outputs were very similar in terms of each metric of species diversity and species abundance (see Figs. B2-B5). As can be seen in Fig. B2, the ODE and SEM yield very similar results in species richness and effective species richness. In some cases, the ODE and SEM differ in their richness output. However, these differences are fairly small. The mean percentage error (MPE) of the ODE approximation, as calculated by

$$\text{MPE} = \frac{1}{S}\sum_{i=1}^{S}\left|\frac{D_{\text{SEM},i} - D_{\text{ODE},i}}{D_{\text{SEM},i}}\right| \times 100 \tag{B.14}$$

where $S$ is the total number of simulations, $D_{\text{SEM},i}$ is the diversity of the $i_{th}$ SEM simulation, and $D_{\text{ODE},i}$ is the diversity the $i_{th}$ ODE model simulation. Over all simulations, MPE = 5.75%. For effective species richness, MPE = 3.64%. Error in which the ODE model predicted greater diversity than the SEM is most likely due to stochastic extinction due to drift. This is particularly likely when diversity is high, where the expected abundance of each species is correspondingly smaller. This conclusion is reinforced by the fact that the ODE more closely approximates effective species richness (which indicates the SEM and ODE model produce highly similar species evenness). Cases in which the ODE model predicted less diversity could be due to several species randomly persisting longer than expected or the transient dynamics not having entirely finished. In addition, some species have growth rates near zero, in which case their persistence depends on chance. Some error also may be due to deterministic differences in the expected behavior of the two models facilitated by the assumptions of the ODE model. However, given that the ODE does not generally over or under-predict the diversity, this effect is probably not very strong. Overall, the outputs strongly indicate that the ODE model captures the expected diversity output of the

SEM.

#### 4 Derivation of invasion criteria

In this section, I derive the approximate invasion criteria of the additive predation model when species experience inter-specific variation in intrinsic ($Y$) with $d = 1$. The per capita growth rate of species $i$, substituting in the seedling abundance values, is

$$\frac{1}{p_i}\frac{dp_i}{dt} = r_i = \delta\Bigg[\frac{Y_i\big[(1-d)+p_i d\big]e^{-a\left(1+p_i E_A\right)}}{Y_i\big[(1-d)+p_i d\big]e^{-a\left(1+p_i E_A\right)} + d\sum_{k\neq i} Y_k p_k e^{-ap_k E_A}} + Y_i d \sum_{m\neq i} \frac{1}{Y_m\big[(1-d)+p_m d\big]e^{-a\left(1+p_m E_A\right)} + d\sum_{k\neq m} Y_k p_k e^{-ap_m E_A}} p_k - 1\Bigg] \tag{B.15}$$

Species $i$ can invade is this quantity if positive when it is rare ($p_i \to 0$). When $d = 1$ (the case of interest), the above reduces to

$$Y_i \sum_{m\neq i} \frac{p_m}{Y_m p_m e^{-a\left(1+p_m E_A\right)} + \sum_{k\neq m} Y_k p_k e^{-ap_k E_A}} > 1 \tag{B.16}$$

To further simplify the above equation, I remove the term $e^{-a\left(1+p_m E_A\right)}$ and substitute it with $e^{-ap_m E_A}$ in the denominator of the summation. Doing so yields

$$Y_i \sum_{m\neq i} \frac{p_m}{\sum_{k\neq i} Y_k p_k e^{-ap_k E_A}} > 1 \tag{B.17}$$

This simplification is equivalent to making the species identity of the tree previously occupying a patch (the tree that dies) irrelevant (i.e., JCEs only result from trees nearby the patch rather than the previous occupant of the patch). As long as predation occurs over a non-trivial distance – that is, so long as $\nu$ is not very small – this assumption should not meaningfully affect the invasion criteria. Numerical simulations show that, for the values considered in the main text (5, 7.5, and 10), this simplification does not qualitatively affect results (see "Validation of invasion criteria").

With this simplification, the denominator of equation (B.17) is no longer directly dependent on $m$ and equation (B.17) can be rewritten as

$$Y_i\left(\sum_{m\neq i} p_m\right)\left(\frac{1}{\sum_{k\neq i} Y_k p_k e^{-ap_k E_A}}\right) > 1 \tag{B.18}$$

Because $\sum_{m\neq i} p_m = 1$, the inequality can be written as

$$Y_i > \sum_{k\neq i} Y_k p_k e^{-ap_k E_A} \tag{B.19}$$

Taking the 1st order Taylor expansion of $p_k e^{-ap_k E_A}$ around the point $1/N$ where $N$ is the number of species in the resident community yields a close approximation of the expression so long as inter-specific variation in $p_k$ is not very. Thus,

$$p_k e^{-ap_k E_A} \approx \frac{e^{-aE_A/N}}{N} - \frac{1}{N}e^{-aE_A/N}\left(1-\frac{aE_A}{N}\right) + p_k e^{-aE_A/N}\left(1-\frac{aE_A}{N}\right) \tag{B.20}$$

Substituting this into the above equation yields

$$Y_i > \sum_{k\neq i} Y_k\left(\frac{e^{-aE_A/N}}{N} - \frac{1}{N}e^{-aE_A/N}\left(1-\frac{aE_A}{N}\right) + p_k e^{-aE_A/N}\left(1-\frac{aE_A}{N}\right)\right) \tag{B.21}$$

Notably, the summation can be broken into two terms:

$$\sum_{k\neq i} Y_k\left(\frac{e^{-aE_A/N}}{N} - \frac{1}{N}e^{-aE_A/N}\left(1-\frac{aE_A}{N}\right)\right) + \sum_{k\neq i} p_k Y_k e^{-aE_A/N}\left(1-\frac{aE_A}{N}\right) \tag{B.22}$$

The first term in the summation is simple:

$$\begin{aligned}\sum_{k\neq i} Y_k\left(\frac{e^{-aE_A/N}}{N} - \frac{1}{N}e^{-aE_A/N}\left(1-\frac{aE_A}{N}\right)\right) &= \frac{1}{N}\sum_{k\neq i} Y_k\left(e^{-aE_A/N} - e^{-aE_A/N}\left(1-\frac{aE_A}{N}\right)\right)\\ &= \overline{Y}ae^{-aE_A/N} - \overline{Y}e^{-aE_A/N}\left(1-\frac{aE_A}{N}\right)\end{aligned} \tag{B.23}$$

where $\overline{Y}$ is the mean intrinsic fitness of the resident community.

The second term in the summation can be expressed by noting the property

$$\frac{1}{N}\sum_{m=1}^{N} A_m B_m = \overline{A}\times\overline{B} + \mathrm{Cov}(A,B) \tag{B.24}$$

Using this property and substituting $A$ and $B$ with $p$ and $Y$ yields

$$\begin{aligned}\sum_{k\neq i} p_k Y_k e^{-aE_A/N}\left(1-\frac{aE_A}{N}\right) &= e^{-aE_A/N}\left(1-\frac{aE_A}{N}\right)N\frac{1}{N}\sum_{k\neq i} p_k Y_k\\ &= e^{-aE_A/N}\left(1-\frac{aE_A}{N}\right)N\left(\frac{\overline{Y}}{N} + \mathrm{Cov}(p,Y)\right)\\ &= \overline{Y}e^{-aE_A/N}\left(1-\frac{aE_A}{N}\right) + e^{-aE_A/N}\left(1-\frac{aE_A}{N}\right)N\mathrm{Cov}(p,Y)\end{aligned} \tag{B.25}$$

Adding equations (B.23) and (B.25) yields

$$\overline{Y}e^{-aE_A/N} + e^{-aE_A/N}\left(1-\frac{aE_A}{N}\right)N\mathrm{Cov}(p,Y) \tag{B.26}$$

Using and implementing this expressions and plugging it into the invasion criteria, species $i$ can invade if

$$Y_i > \overline{Y}e^{-aE_A/N} + e^{-aE_A/N}\left(1-\frac{aE_A}{N}\right)N\mathrm{Cov}(p,Y) \tag{B.27}$$

which can be rearranged as

$$Y_i > \underbrace{\overline{Y}}_{\substack{\text{mean}\\ \text{fitness}}} \underbrace{e^{-a\frac{E_A}{N}}}_{\text{mean JCE}} \left[1 + \underbrace{\frac{\mathrm{Cov}(p, Y)}{\overline{Y}/N}}_{\substack{\text{covariance}\\ \text{term}}} \underbrace{\left(1 - a\frac{E_A}{N}\right)}_{\substack{\text{covariance}\\ \text{coefficient}}}\right] \tag{B.28}$$

which is identical to equation (7) from the main text.

#### 5 Figures

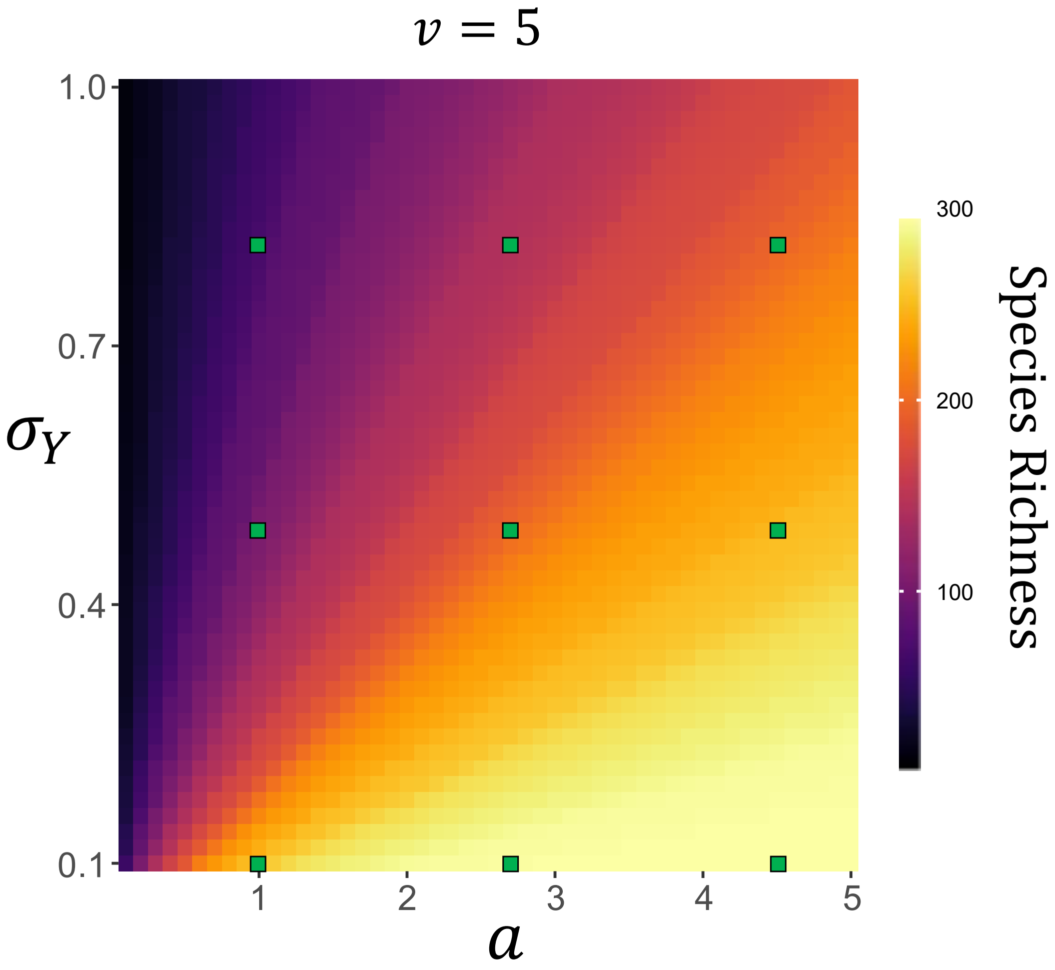

**Fig. B1** Visualization of the parameter values used for the SEM. Green squares depict the parameter values run in each SEM simulation. Values are superimposed on the heatmaps from the main text to show the extent of parameter space examined. The left hand figure depicts when species vary in intrinsic fitness and the right hand figure depicts when species vary in JCE susceptibility. $\nu = 5$ is noted on each figure for consistency with the figures from the main text, but the same parameters were tested for the $\nu = 7.5$ and $\nu = 10$ cases as well.

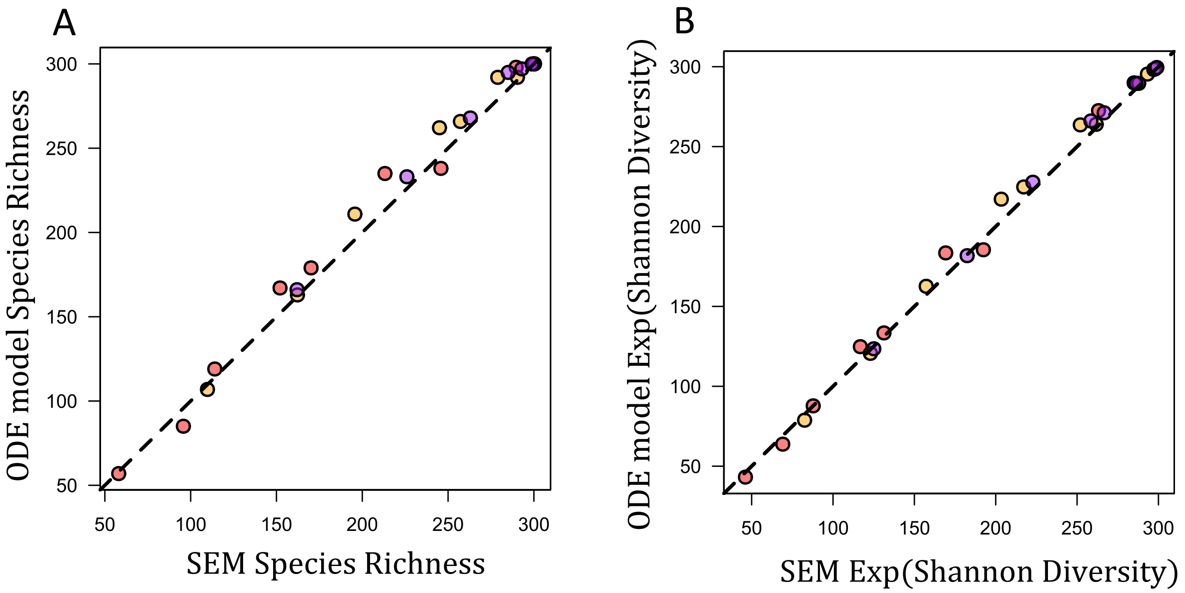

**Fig. B2** ODE model validation. The figures compare species richness and effective species richness (the exponential of Shannon Diversity) between SEM and ODE model simulations under identical parameterizations. The dashed line is the one-to-one line (points on the line represent when the SEM and ODE yield the exact same diversity output). Red points are when $\nu = 5$, orange/yellow points are when $\nu = 7.5$, and purple points are when $\nu = 10$. To a first approximation, the ODE model yields the same output as the SEM in both diversity metrics.

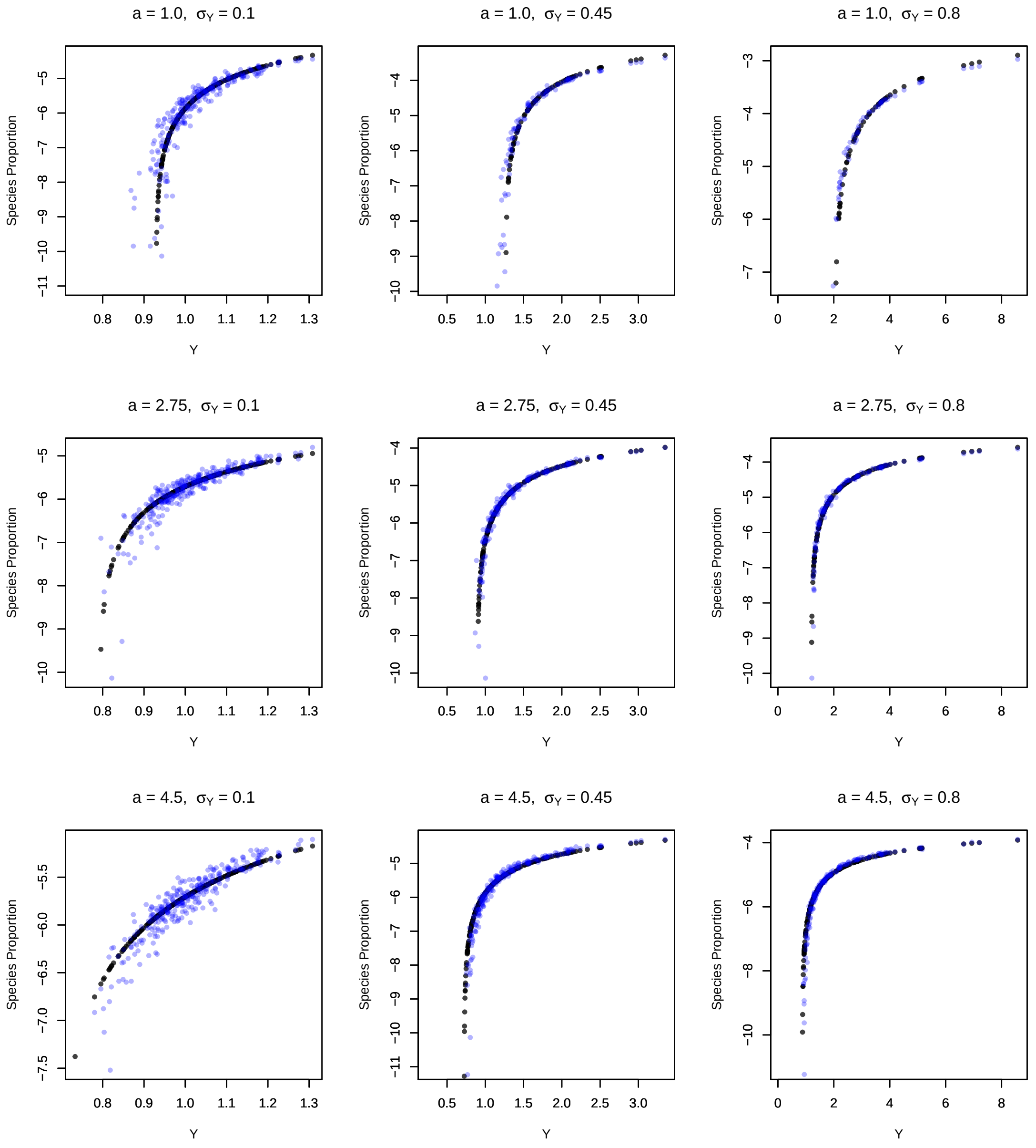

**Fig. B3** Comparisons between identical parameterizations of the ODE approximation (black) and SEM (blue) outputs under nine parameter values when species vary in intrinsic fitness ($Y$). The$y$-axis depicts the log-proportion of each species and the $x$-axis depicts $Y$ of each species. These parameter values span the most of the parameter space explored in Fig. 4 of the main text. In all plots, $\nu = 5$, $\gamma = 0.2$, and $d = 1.0$. Other relevant parameters are listed on each plot.

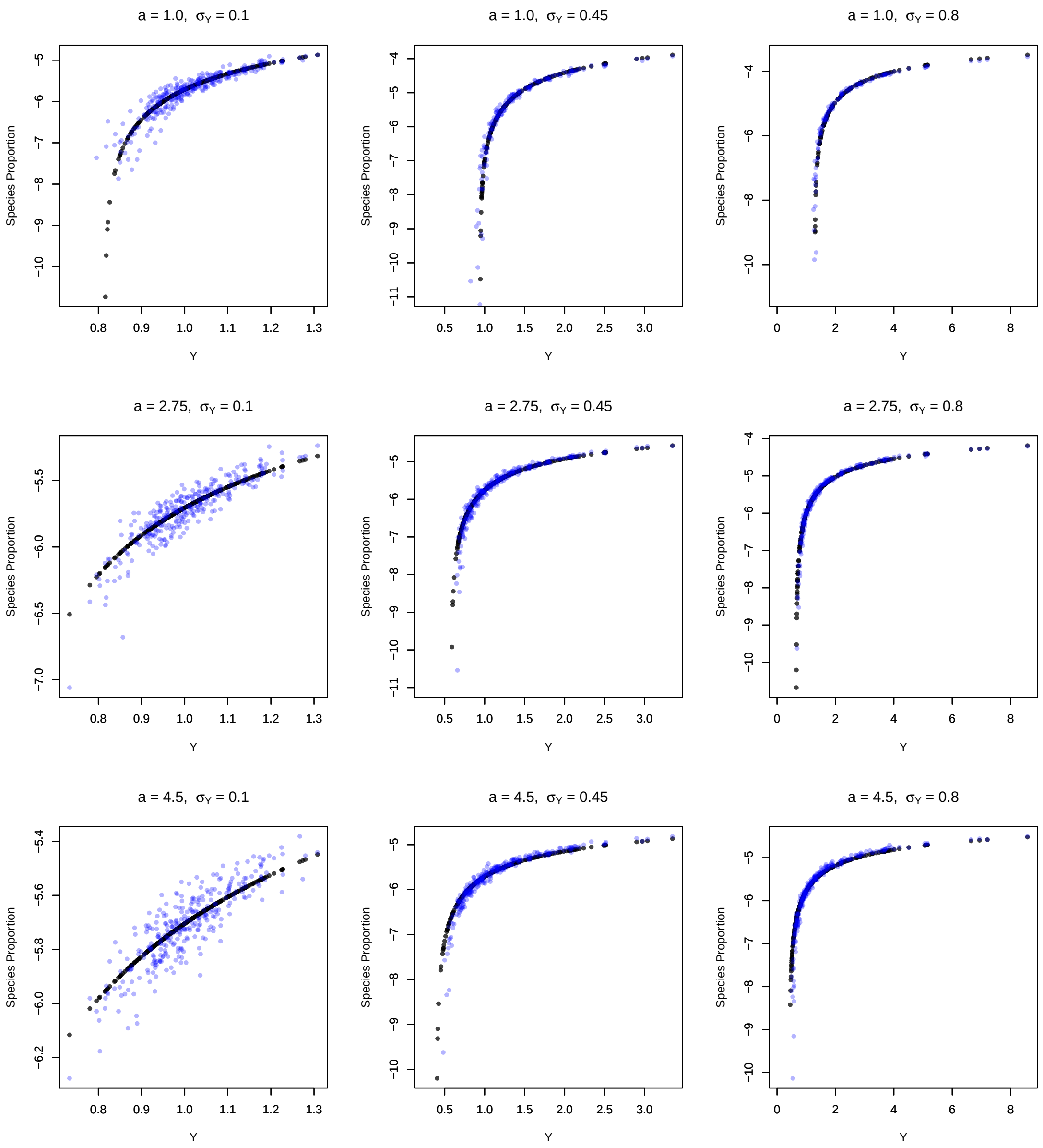

**Fig. B4** The same format as Fig. B3, but with $\nu = 7.5$.

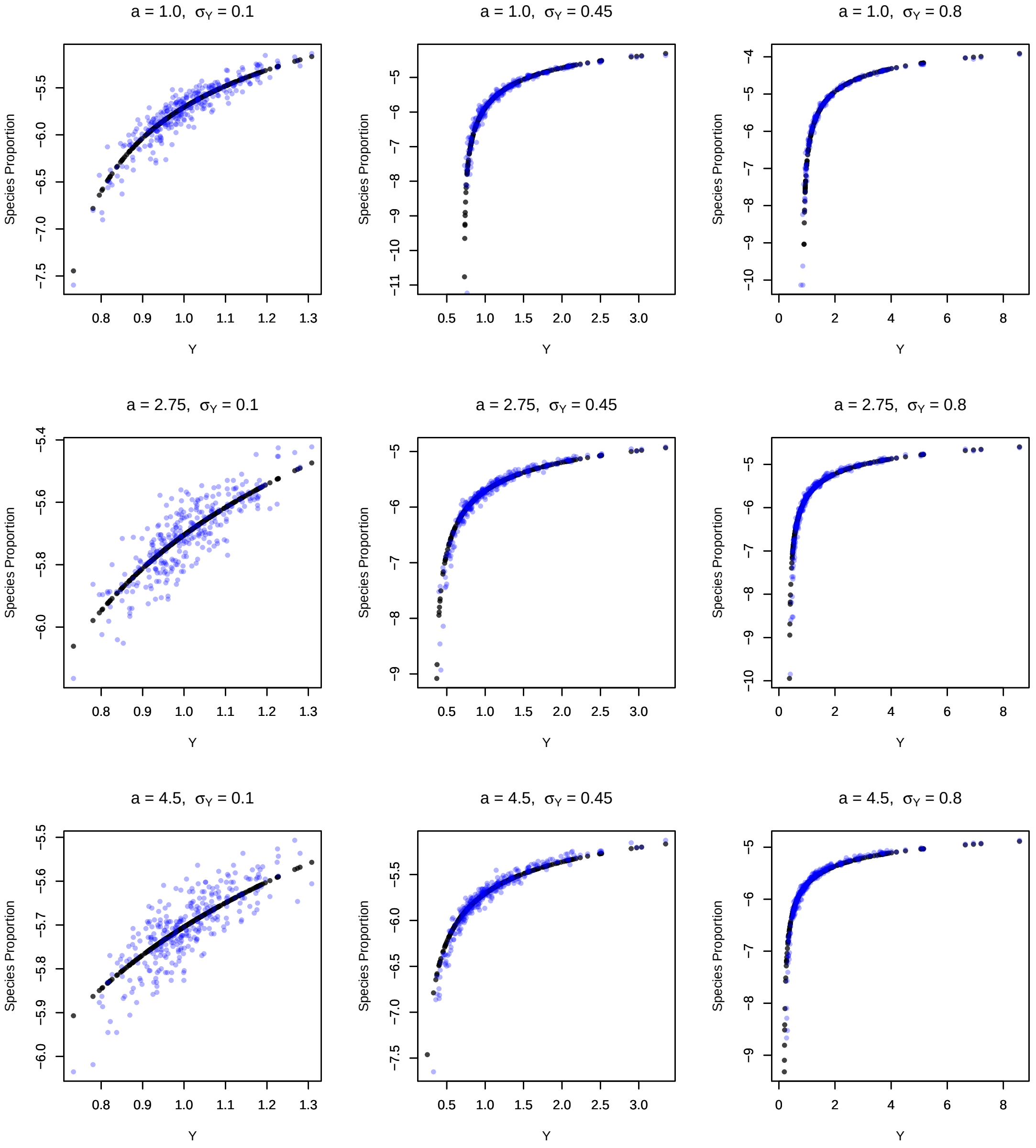

**Fig. B5** The same format as Fig. B3, but with $\nu = 10$.

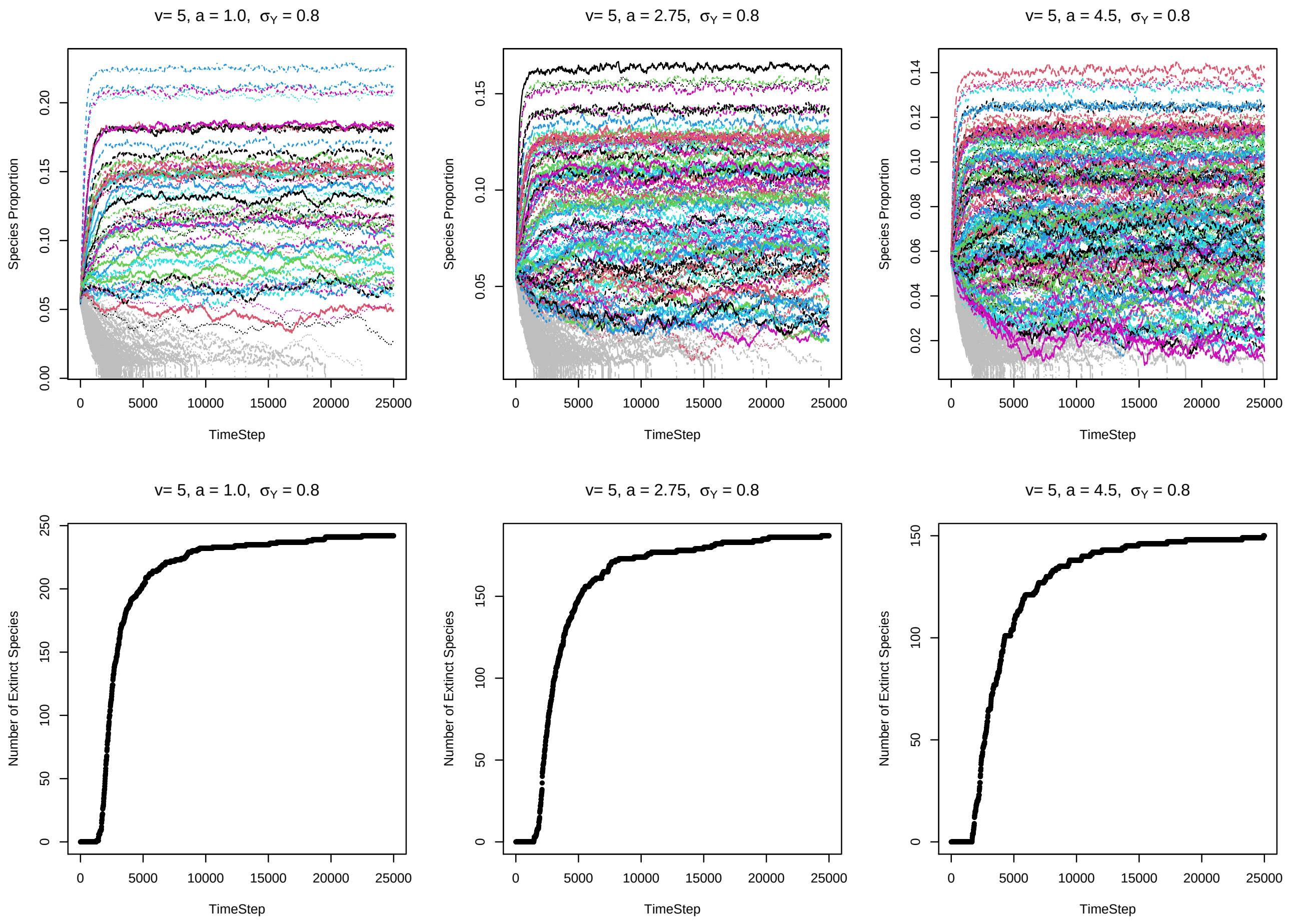

**Fig. B6** Examples of the SEM simulation time series outputs. The top row shows examples of the time series outputs of the SEMs. The $x$-axis is time and the $y$-axis is each species' proportion. Proportions have been square-root transformed to aid visualization. Colored trajectories indicate species that persisted throughout the simulation; grey trajectories indicate species that went extinct. In these examples, $Y$ varies between species. Parameters are listed on each plot. Dynamics as shown are typical examples from the SEMs. Most species settle into a relatively stable pseudo-equilibrium, while lower abundance species fluctuate due to drift. The bottom row shows the number of extinct species in the community as a function of time. Each panel corresponds to the plot above it. As can be seen, most species that go extinct do so in the early stages of the dynamics. Therefore, the vast majority of persisting species likely persist deterministically.

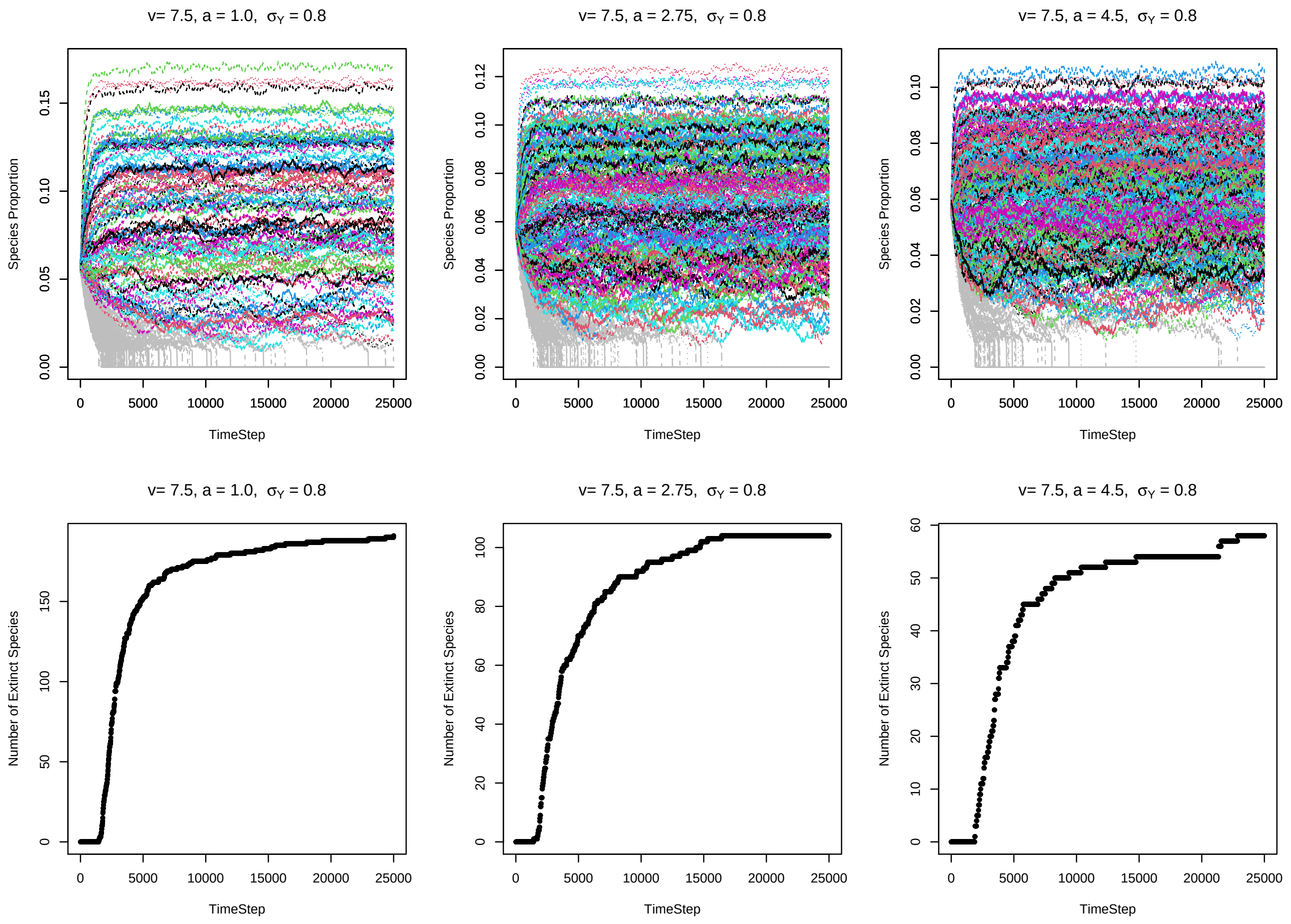

**Fig. B7** The same as Fig. B7, but with $\nu = 7.5$.

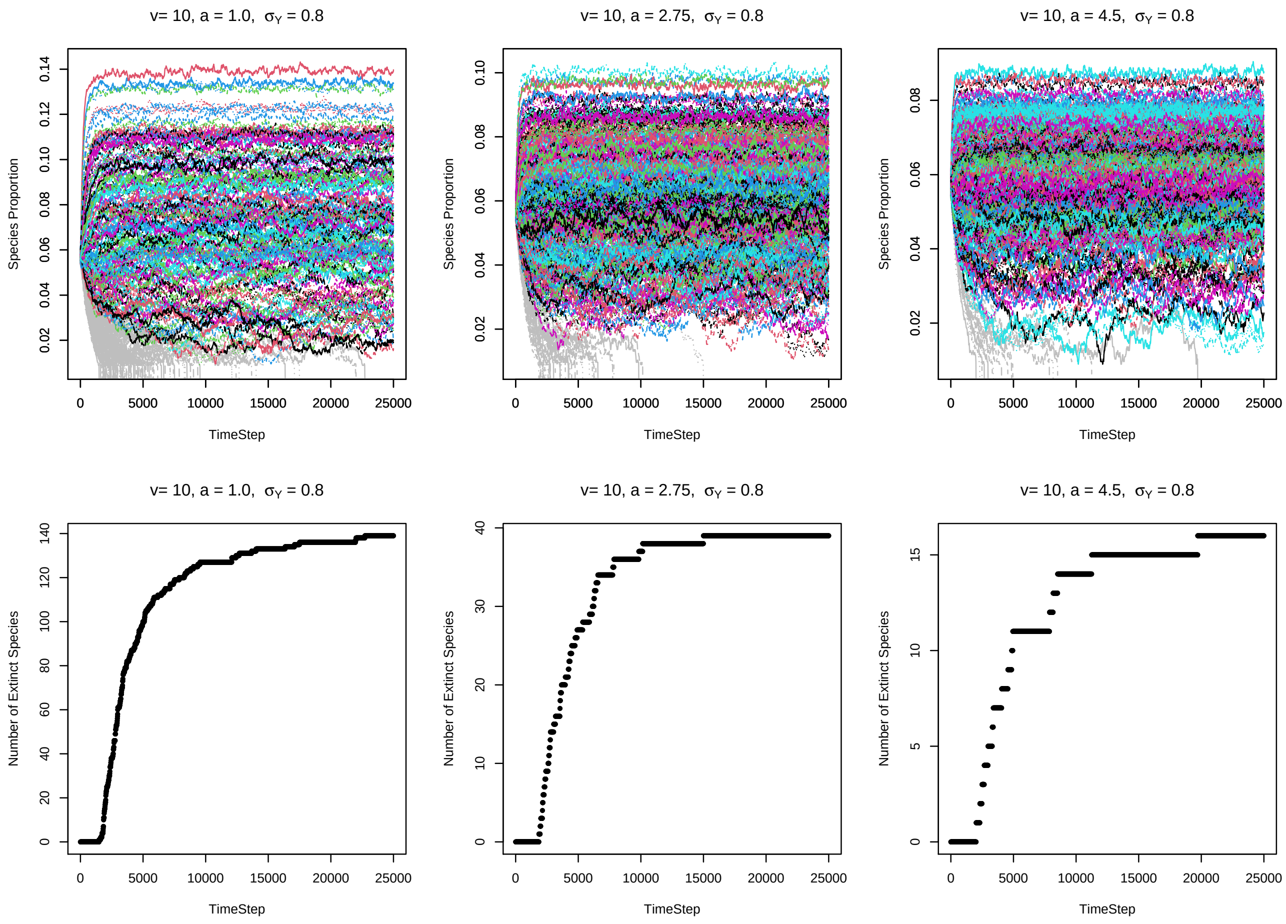

**Fig. B8** The same as Fig. B7, but with $\nu = 10$.
