## Appendix C for "The functional form of specialized predation dramatically affects whether Janzen-Connell effects can prevent competitive exclusion"

### Appendix C: Invasion criteria validation

July 10, 2021

### 1 Introduction

In this appendix, I validate the invasion criteria approximation of each model (the fixed neighborhood model and the additive predation model). See Appendices A and B for the derivation of the approximations. This appendix is written assuming the reader is familiar with the invasion criteria presented in the main text and the derivations in Appendices A and B.

First, I show each invasion criteria approximation can be compared to a simple expression derived from the ODE expressions. Second, I use these expressions to derive exact and approximate values of $\Phi$, the metric used in the main text to compare the invasion criteria of each Janzen Connell Effect (JCE) functional form. Third, I show numerical results comparing the exact and approximate values of $\Phi$.

### 2 Derivation of comparisons

#### 2.1 Additive Predation Model

As shown in the main text and Appendix B, the invasion criteria of the additive predation model is approximately:

$$Y_i > \underbrace{\overline{Y}}_{\substack{\text{mean}\\\text{fitness}}} \underbrace{e^{-a\frac{E_A}{N}}}_{\text{mean JCE}} \left[1 + \underbrace{\frac{\mathrm{Cov}(p,Y)}{\overline{Y}/N}}_{\substack{\text{covariance}\\\text{term}}} \underbrace{\left(1 - a\frac{E_A}{N}\right)}_{\substack{\text{covariance}\\\text{coefficient}}}\right] \tag{C.1}$$

This equation is approximated from the following expression:

$$Y_i \sum_{m\neq i} \frac{p_m}{Y_m p_m e^{-a\left(1+p_m E_A\right)} + \sum_{k\neq m} Y_k p_k e^{-a p_k E_A}} > 1 \tag{C.2}$$

This is shown in Appendix B. The expression can be rearranged as

$$Y_i > \frac{1}{\sum_{m\neq i} \frac{p_m}{Y_m p_m e^{-a\left(1+p_m E_A\right)} + \sum_{k\neq m} Y_k p_k e^{-a p_k E_A}}} \tag{C.3}$$

Equations (C.3) and (C.1) have the same value of the left hand side ($Y_i$) and are thus analogous.

#### 2.2 Fixed Neighborhood Model

As shown in the main text and Appendix A, the invasion criteria of the fixed neighborhood model is approximately:

$$Y_i > \underbrace{\overline{Y}}_{\substack{\text{mean}\\ \text{fitness}}} \underbrace{e^{-\left(1-e^{-a}\right)\frac{E_n}{N}}}_{\text{mean JCE}} \left[1 + \underbrace{\frac{\text{Cov}(p,Y)}{\overline{Y}/N}}_{\substack{\text{covariance}\\ \text{term}}} \underbrace{\left(1 - \left(1 - e^{-a}\right)\frac{E_n}{N}\right)}_{\substack{\text{covariance}\\ \text{coefficient}}}\right] \tag{C.4}$$

This approximation is derived from the below expression:

$$Y_i \sum_{m \neq i} \frac{p_m}{Y_m e^{-a} p_m + \sum_{k \neq m} Y_k p_k \left(e^{-p_k E_n} + e^{-a}(1 - e^{-p_k E_n})\right)} > 1 \tag{C.5}$$

This is shown in detail in Appendix A. The expression can be rearranged as

$$Y_i > \frac{1}{\sum_{m \neq i} \frac{p_m}{Y_m e^{-a} p_m + \sum_{k \neq m} Y_k p_k \left(e^{-p_k E_n} + e^{-a}(1 - e^{-p_k E_n})\right)}} \tag{C.6}$$

Akin to the additive predation model, equations (C.6) and (C.4) have the same value of the left hand side ($Y_i$) and are thus analogous.

### 3 Comparison of $\Phi$

The main text examines the relative ability of each model to promote coexistence using the metric $\Phi$ (main text, equation (9)). $\Phi$ is equal to equation (C.4) divided by equation (C.1), or

$$\Phi = \underbrace{e^{\frac{\gamma\pi}{N}\left(2a\nu^2 - (1-e^{-a})r^2\right)}}_{\text{mean JCE ratio}} \underbrace{\frac{1 + \frac{\text{Cov}(p,Y)}{\overline{Y}/N}\left(1 - \left(1 - e^{-a}\right)\frac{\pi\gamma r^2}{N}\right)}{1 + \frac{\text{Cov}(p,Y)}{\overline{Y}/N}\left(1 - a\frac{2\pi\gamma\nu^2}{N}\right)}}_{\text{covariance ratio}} \tag{C.7}$$

Therefore, to validate the invasion criteria approximations as used in the main text, I compare the above approximate value of $\Phi$ to its exact value under identical parameterizations. However, it is first necessary to modify $\Phi$. $\Phi$ in the main text assumes that $N$ and $\text{Cov}(p, Y)$ are exactly for the same in each model for a given parameterization. In practice, this is almost certainly untrue. Therefore, it is necessary to substitute the values for each model. This yields

$$\Phi_{approx} = \frac{\overline{Y_n}}{\overline{Y_A}} \frac{e^{-\left(1-e^{-a}\right)\frac{\pi\gamma r^2}{N_n}}}{e^{-a\frac{2\pi\gamma\nu^2}{N_A}}} \frac{1 + \frac{\text{Cov}_n(p,Y)}{\overline{Y_n}/N_n}\left(1 - \left(1 - e^{-a}\right)\frac{\pi\gamma r^2}{N_n}\right)}{1 + \frac{\text{Cov}_A(p,Y)}{\overline{Y_A}/N_n}\left(1 - a\frac{2\pi\gamma\nu^2}{N_A}\right)} \tag{C.8}$$

where, given a parameterization, $N_n$ is the number of species in the community for the fixed neighborhood model, $\text{Cov}_n(p, Y)$ is the covariance between intrinsic fitness and abundance for the fixed neighborhood model, $\overline{Y_n}$ is the mean fitness of the resident community for the fixed neighborhood model, $N_A$ is the number of species in the community for the additive predation model, $\text{Cov}_A(p, Y)$ is the covariance between intrinsic fitness and abundance for the additive predation model, and $\overline{Y_A}$ is the mean fitness of the resident community for the additive predation model.

The exact value of $\Phi$ is the quotient of equation (C.6) and equation (C.3):

$$\Phi_{exact} = \frac{\sum_{m\neq i} \frac{p_m}{Y_m e^{-a} p_m + \sum_{k\neq m} Y_k p_k (e^{-p_k E_n} + e^{-a}(1 - e^{-p_k E_n}))}}{\sum_{m\neq i} \frac{p_m}{Y_m p_m e^{-a(1+p_m E_A)} + \sum_{k\neq m} Y_k p_k e^{-a p_k E_A}}} \tag{C.9}$$

To examine the quality of $\Phi$ as an approximation, I compare $\Phi_{approx}$ to $\Phi_{exact}$.

#### 4 Numerical comparisons

##### 4.1 Parameterizations

I conducted a number of numerical computations of $\Phi_{approx}$ and $\Phi_{exact}$. For each model, I examined the quality of the approximation for a range $a$ values ranged between $1.0 - 5.0$ under several conditions. I considered resident communities starting with $N = 300$ species under the condition that $Y \sim \text{lognormal}[\mu = 0, \sigma_Y]$ for $\sigma_Y = 0.1, 0.25$, and 0.5. I use this range of $\sigma_Y$ (lower than the range of the main paper, $0.1 - 1.0$) because the fixed neighborhood model yields very low diversity for higher $\sigma_Y$ and the approximation assumes that $N$ is somehwat large. For each model, I examined $\nu = r = 5$, 7.5, and 10. ODE simulations were run for 6,250 generations, more than sufficient for the system to reach equilibrium. I considered species $i$ extinct if $\log(p_i) < -11$, noting that $\log(p_i) = -11$ corresponds to a species with an expected abundance of approximately 1 tree at BCI ($\log(1/86000) \approx -11.4$). After the simulations were run, I numerically calculated $\Phi_{approx}$ and $\Phi_{exact}$. Simulations were performed in R (R Core Team, 2020).

#### 4.2 Results

Fig. C1 shows the general quality of $\Phi_{approx}$ compared to $\Phi_{exact}$. For the majority of cases, the approximation yields similar outputs. For higher values of $\Phi$, $\Phi_{approx}$ tends to slightly underestimate $\Phi_{exact}$. The higher $\Phi$ is, the more strongly the additive predation model reduces fitness differences. Therefore, the approximation tends to slightly underestimate the stabilizing effects of the additive predation model relative to the fixed neighborhood model.

Notably, the range of $\Phi$ is much smaller than that of the main text (the range on the axes of Fig. C1 to the range on the y-axes in Fig. 3 from the main text). This reflects the difference in diversity maintained by each model. For each invasion criteria, $N$ (the number of species in the resident community) strong affects the invasion criteria. Under identical parameterizations, the fixed neighborhood model maintains considerably less diversity – thus, $N_n << N_A$ in most cases, yielding a lower value of $\Phi$. However, overall, it is clear that $\Phi_{approx} \approx \Phi_{exact}$.

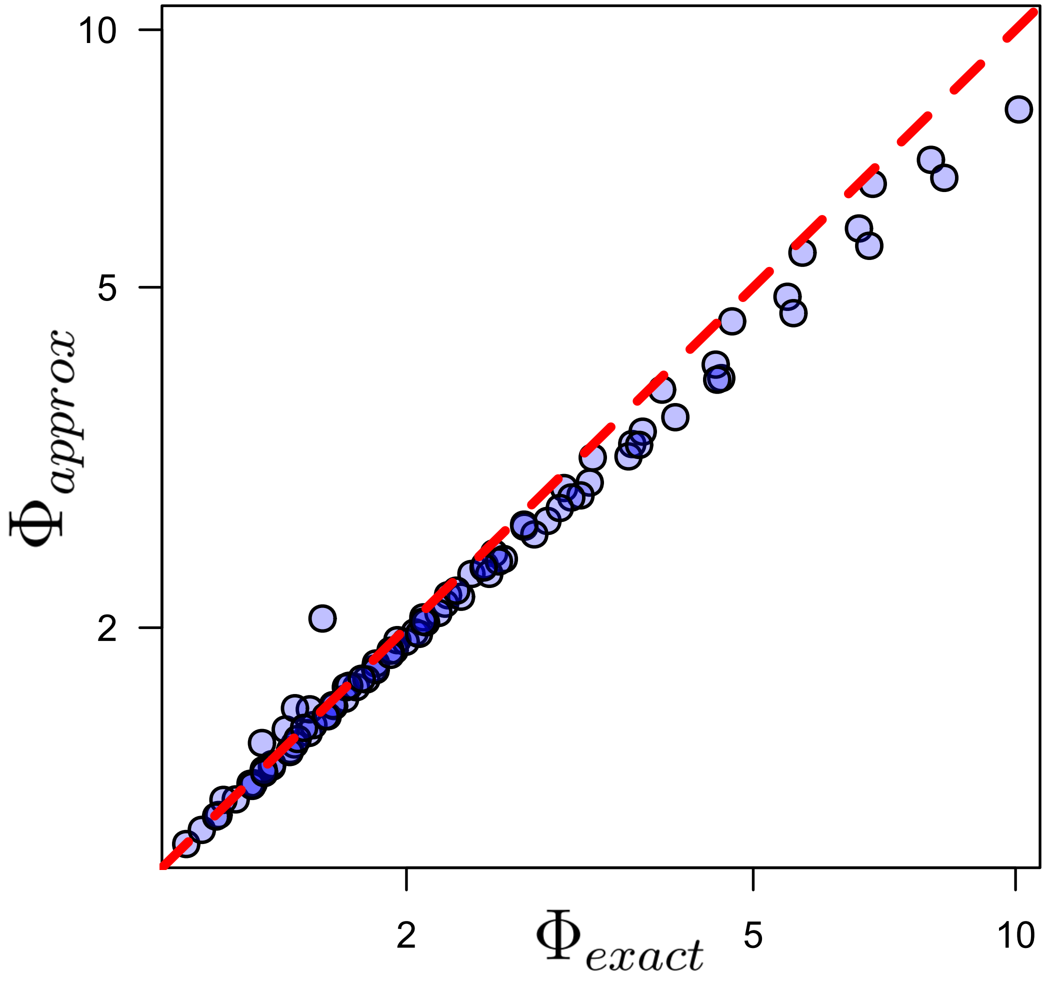

**Fig. C1** Comparison $\Phi_{approx}$ and $\Phi_{exact}$ across all parameterizations noted in the text of this appendix. Note that both axes are log-scaled. The dashed line is the one-to-one line (points on the line represent when the approximation and exact value perfectly coincide). In general, the approximation and exact values yield highly similar outputs.
