## Appendix D for "The functional form of specialized predation dramatically affects whether Janzen-Connell effects can prevent competitive exclusion"

### Appendix D: Nearest neighbors of common conspecific adults for the additive predation model

July 10, 2021

#### Contents

**1 Introduction** **2**

**2 Nearest-Neighbors** **2**

2.1 Distance calculation . . . . . . . . . . . . . . . . . . . . . . . . . . . . . . . 2

2.2 Results . . . . . . . . . . . . . . . . . . . . . . . . . . . . . . . . . . . . . . . 3

**3 Tables** **5**

### 1 Introduction

Previous studies note that conspecific trees of common species in tropical forests are somewhat close in space (on the order of 20 meters apart; Hubbell (1980)). Under the fixed neighborhood model, as Chisholm & Fung (2020) note, this spacing inhibits the feasible neighborhood effect size ($E_n$) when JCEs are strong. Chisholm & Fung (2020) argue that JCEs must be either weak or $E_n$ must be very small. In this appendix, I briefly examine the impact Janzen-Connell Effects (JCEs) on the minimum distance between conspecific adults under the additive predation model. I examine the minimum distance between conspecific adults based on the Spatially Explicit Model (SEM) simulations described in Appendix B. I show that the average distance between conspecific adults of the most common species in the community is almost always less than 20 meters apart.

### 2 Nearest-Neighbors

#### 2.1 Distance calculation

Each SEM simulation was conducted on a $275 \times 275$ matrix. Each entry of the matrix contains an integer, $i$, between 1 and 300 that depicts an individual tree of species $i$. The location of each individual tree can be represented through coordinates of the matrix. Let $(x, y)$ be a given patch of the matrix where $x$ corresponds to a column of the matrix and $y$ corresponds to a row of the matrix.

To calculate the distance between conspecifics, I calculated the euclidean distances between center points of patches. The SEM was constructed with an average density of $\gamma = 0.20$ individuals per meter, in which case distance between patches is $1/\sqrt{\gamma}$ meters. Given the location of an individual tree of species $i$ at location $(x, y)$, the distance to its nearest conspecific neighbor is:

$$D^i_{x,y} = \frac{1}{\sqrt{\gamma}} \min\left(\sqrt{(x - \overrightarrow{x_i})^2 + (y - \overrightarrow{y_i})^2}\right) \tag{D.1}$$

where $D^i_{x,y}$ is the minimum distance between species $i$ at position $(x, y)$ and a conspecific adult,

$\overrightarrow{x_i}$ is a vector of all column positions of individuals of species $i$ and $\overrightarrow{y_i}$ is a vector of all column positions of individuals of species $i$. These vectors exclude the individual at $(x, y)$. This equation is slightly modified in its actual implementation because the simulations were conducted on a torus. For all SEM simulations of additive predation model (see Appendix B for details) I calculated the minimum distance between conspecifics of all adults of the most common species in the community ($D^i_{x,y}$ calculated for all $\overrightarrow{x_i}$ and $\overrightarrow{y_i}$). From the resulting distribution of minimum distances, I calculated the mean distance of the most common species to its nearest conspecific neighbor, $\overline{D^i_{x,y}}$.

#### 2.2 Results

$\overline{D^i_{x,y}}$ was below 20 meters in all but one case (when $\sigma_Y = 0.10$, $a = 4.5$, and $\nu = 10$). Tables C1-C3 list all the results.

$\overline{D^i_{x,y}}$ tended to be lowest when $\sigma_Y$ was high and $a$ was low. This reflects two factors. Higher $\sigma_Y$ increases the relative abundance of the most common species (via an increase in its relative fitness). The more common a species in the community is, the shorter the average distance between conspecific neighbors. Higher $a$ corresponds to higher $\overline{D^i_{x,y}}$ because $a$ defines the strength of JCEs. As JCEs make the local environment less hospitable for conspecific seedlings and can influence the spatial distribution of species to be more uniformly distributed in space than randomly expected (Hubbell, 1980; Levi *et al.*, 2019) at least at smaller spatial scales (Detto & Muller-Landau, 2013). Therefore, larger $a$ tends to increase $\overline{D^i_{x,y}}$. $\overline{D^i_{x,y}}$ also increased with higher $\nu$. This is unsurprising: the lager $\nu$ is, the larger the spatial scale of specialized predation.

#### 3 Tables

| **Parameters** | $a = 1.0$ | $a = 2.75$ | $a = 4.5$ |
|---|---|---|---|
| $\sigma_Y = 0.10$ | 10.96 | 13.97 | 16.58 |
| $\sigma_Y = 0.45$ | 6.69 | 9.78 | 11.76 |
| $\sigma_Y = 0.80$ | 5.51 | 8.22 | 9.84 |

**Table C1:**$\overline{D^i_{x,y}}$ in meters of the most common species. Species vary in $Y$. $d = 1$, $\gamma = 0.20$, and $\nu = 5$. All other parameters are listed on the plot.

| **Parameters** | $a = 1.0$ | $a = 2.75$ | $a = 4.5$ |
|---|---|---|---|
| $\sigma_Y = 0.10$ | 14.00 | 18.06 | 19.77 |
| $\sigma_Y = 0.45$ | 8.98 | 13.15 | 15.11 |
| $\sigma_Y = 0.80$ | 7.41 | 10.95 | 13.31 |

**Table C2:** $\overline{D^i_{x,y}}$ in meters of the most common species. Species vary in $Y$. $d = 1$, $\gamma = 0.20$, and $\nu = 7.5$. All other parameters are listed on the plot.

| **Parameters** | $a = 1.0$ | $a = 2.75$ | $a = 4.5$ |
|---|---|---|---|
| $\sigma_Y = 0.10$ | 16.17 | 19.36 | 21.06 |
| $\sigma_Y = 0.45$ | 10.93 | 15.40 | 18.03 |
| $\sigma_Y = 0.80$ | 8.91 | 13.53 | 16.08 |

**Table C2:** $\overline{D^i_{x,y}}$ in meters of the most common species. Species vary in $Y$. $d = 1$, $\gamma = 0.20$, and $\nu = 10$. All other parameters are listed on the plot.
