## Appendix E for "The functional form of specialized predation dramatically affects whether Janzen-Connell effects can prevent competitive exclusion"

### Appendix E: Supplemental figures

July 10, 2021

#### Contents

1 Introduction 2

2 Simulation notes 2

3 Figures 2

### 1 Introduction

This Appendix contains the supplemental figures referenced in the main text.

### 2 Simulation notes

All of the figures depict outputs from the ODE model. In all simulations, I assumed $\gamma = 0.20$ (near the lower bound at BCI) unless otherwise noted. The initial number of species in the community ($N$) was set to 300 on the basis that BCI contains approximately 300 woody plant species (Condit *et al.*, 2019). ODE simulations were run for 6250 generations, sufficient for the system to reach equilibrium. I considered species $i$ extinct if $\log(p_i) < -11$, noting that $\log(p_i) = -11$ corresponds to a species with an expected abundance of approximately 1 tree at BCI ($\log(1/86000) \approx -11.4$). Simulations were performed in R (R Core Team, 2020).

### 3 Figures

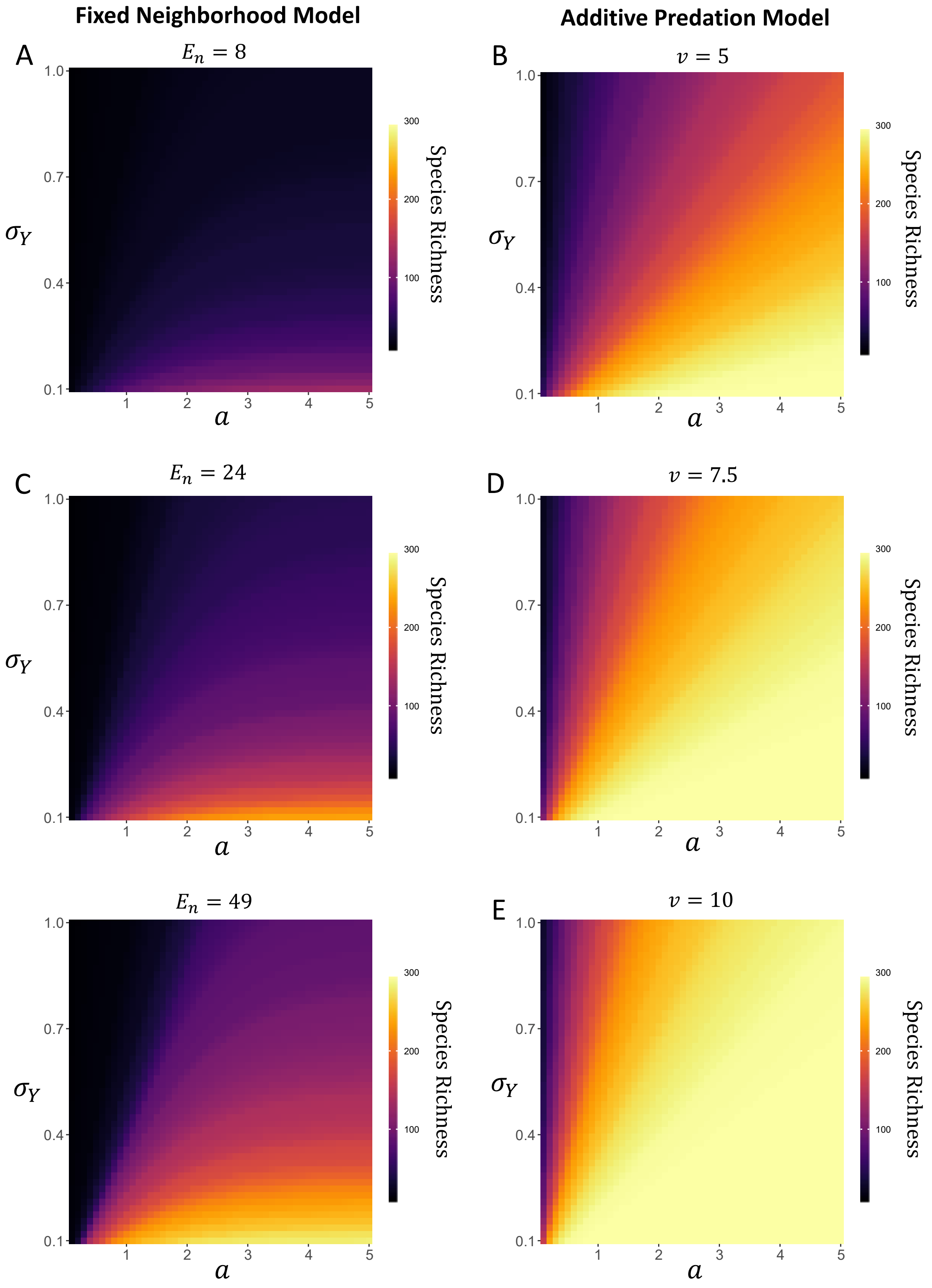


**Fig. D1:** The same as Fig. 4 in the main text, but with $d = 0.6$ instead of $d = 1.0$. While diversity decreases a small amount relative to Fig. 4 from the main text, results are trivially different.

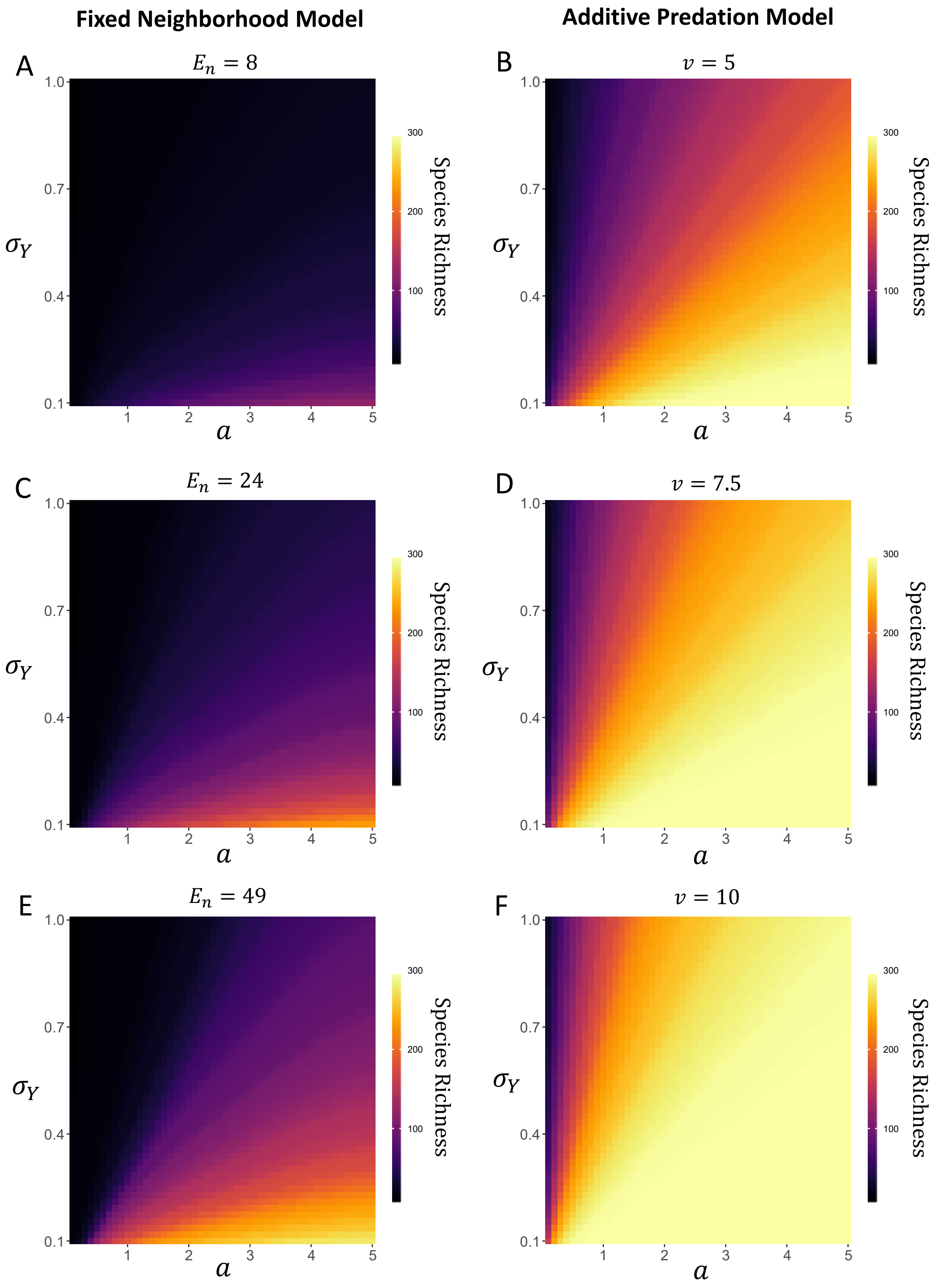


**Fig. D2:** The same as Fig. 4 in the main text, but with $d = 0.1$ instead of $d = 1.0$. Diversity decreases a small amount relative to Fig. 4 from the main text (especially for the fixed neighborhood model) but only a small amount.

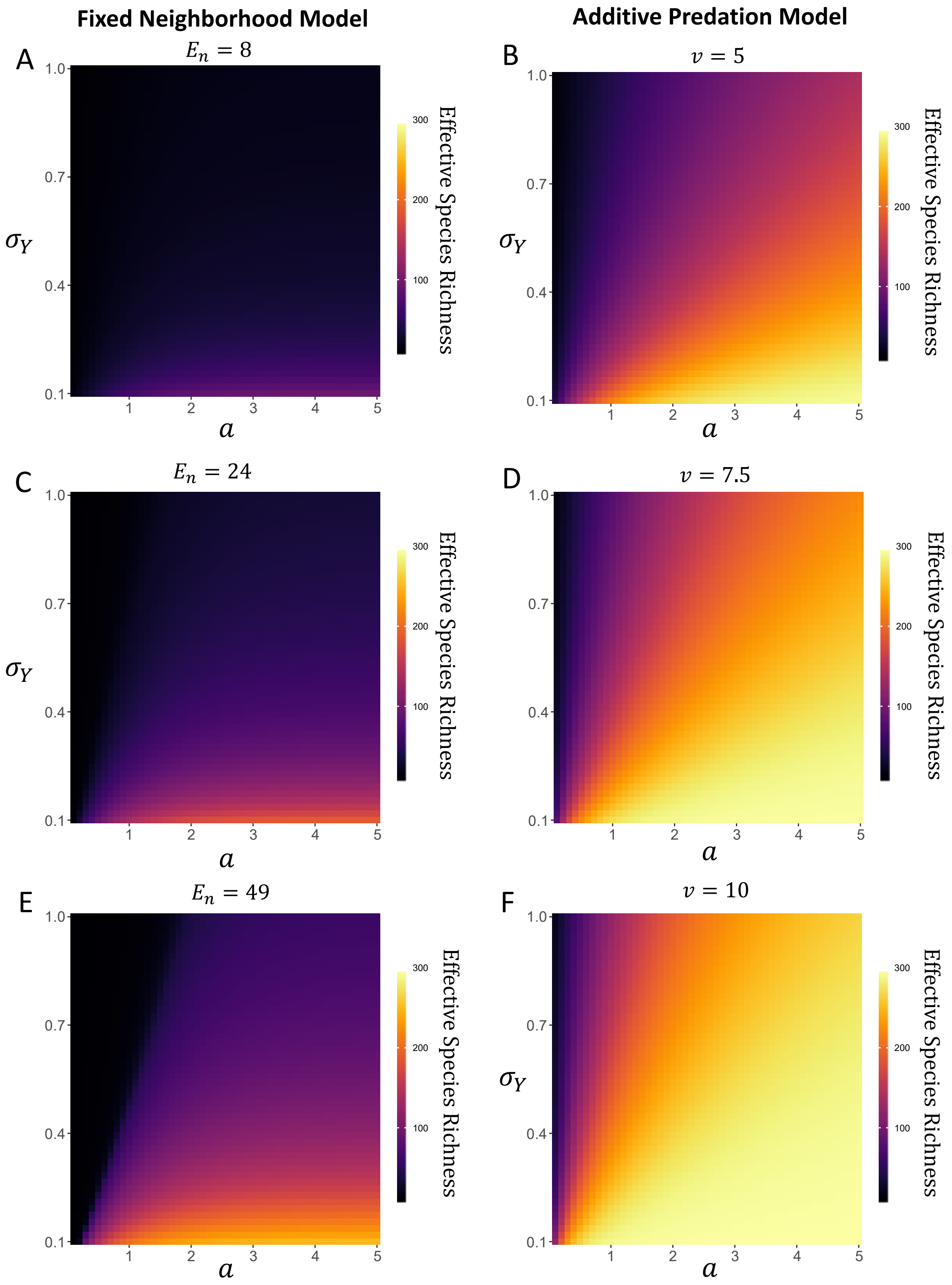


**Fig. D3:** The same as Fig. 4 in the main text, but the $z$-axis shows effective species richness (exp(Shannon Diversity)) instead of species richness. The qualitative patterns are unchanged.

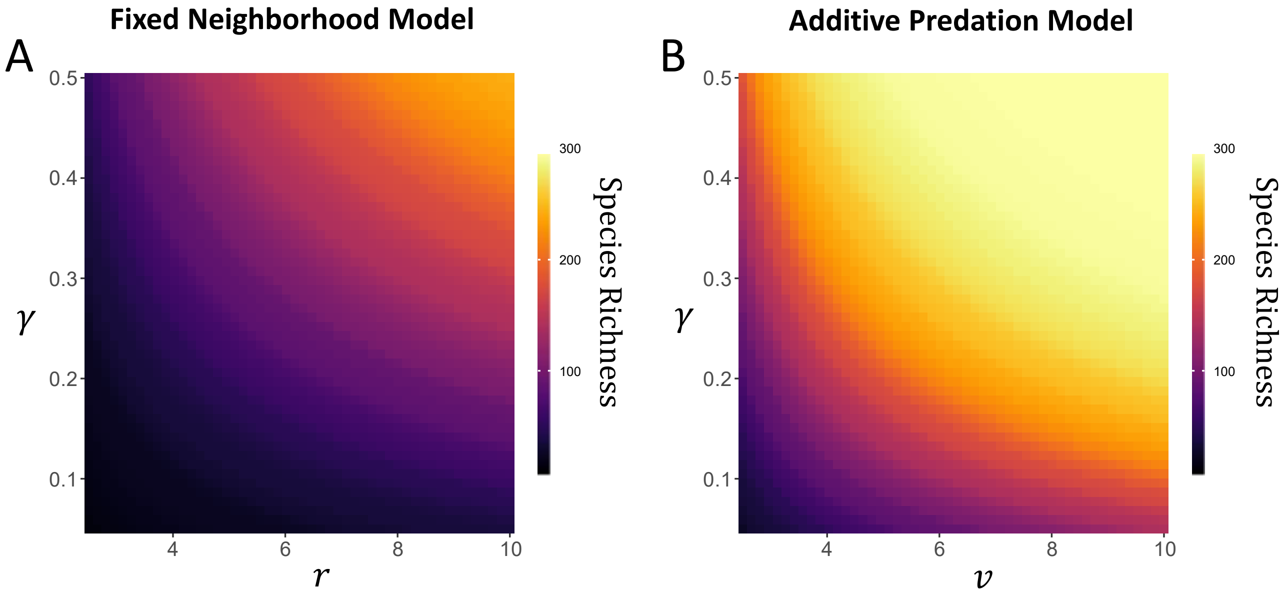


**Fig. D4:** The effect of $\gamma$ and distance-dependence metrics ($r$ and $\nu$) for each JCE functional form. Simulations examine $r$ and $\nu$ between $2.5-10$ and $\gamma$ between $0.05-0.50$. This corresponds to a range of $E_n$ approximately between $1-157$ and $E_A$ between $2-314$. $E_A$ is twice $E_n$ because $E_A = 2\gamma\pi\nu^2 = 2E_n$ if $r = \nu$ given that $E_n = \gamma\pi r^2$. The additive predation model tends to maintain considerably greater diversity. Other parameters are as follows: $d = 1$, $a = 2.5$, and $Y \sim \text{lognormal}[\mu = 0, \sigma_Y = 0.50]$.
